# Quantifying the fitness contribution of phagocytosis

**DOI:** 10.1101/2025.10.23.684244

**Authors:** Paul E. Schavemaker, Kaitlyn S. Tam, Michael Lynch

**Affiliations:** Biodesign Center for Mechanisms of Evolution, Arizona State University, Tempe, AZ, 85287, USA; School of Life Sciences, Arizona State University, Tempe, AZ, 85287, USA

## Abstract

Phagocytosis, the ingestion of cells by other cells, is ubiquitous among eukaryotic life. It is required for food uptake in many single-celled species and for the immune response in multicellular species. The origin of phagocytosis and its role in the evolution of endomembranes and the eukaryotic cell remains obscure. Drawing on a wealth of empirical data, we integrate prey capture, engulfment, and internal and external digestion into a mathematical evolutionary model that quantifies the fitness of a primitive phagocytoser relative to a non-phagocytosing ancestor. We reveal the conditions under which a non-phagocytosing predator that digests its prey externally can persist. We also show that the phagocytoser can outperform the ancestor in a broad range of situations, despite the cost associated with producing a phagocytic cup. Parameter variations delineate how fast engulfment needs to be for phagocytosis to be advantageous, providing clear benchmarks for interpreting the importance of results in genetic knockout studies and mechanical models. The phagocytoser still outperforms the ancestor when food vacuoles can’t fuse back to the plasma membrane, providing arguments in favor of the gradual evolution of phagocytosis and for phagocytosis as the initiator of the endomembrane system.

## Introduction

Cells ingesting other cells, or phagocytosis, is a horrifying spectacle on the microscale in which the life of an organism is snuffed out by being completely engulfed, bathed in acid and hydrolytic enzymes, and finally assimilated into the predator cell. It is a crucial mechanism for the disposal of pathogens in the mammalian immune system^1,2^ as well as the primary mechanism for obtaining sustenance in many single-celled species. The role of phagocytosis in the immune system may have evolved from nutritionally important enteric phagocytes^3^. Observed^4^ as far back as the year 1777, the deeper evolutionary roots of phagocytosis lie in the engulfment of prey cells by predator cells in the microbial world, with many representatives of this mode of sustenance extant across present-day unicellular eukaryotes^5,6^. Phagocytosis occurs in close relatives of the animals, the unicellular choanoflagellates^7^, as well as the more distantly related amoebae^8–11^. Phagocytic representatives from other parts of the eukaryotic tree are present in the heliozoa^12^, dinoflagellates^13,14^, green algae^15,16^, haptophytes^17^, euglenids^18^, and metamonads^19,20^. The most spectacular forms of phagocytosis are found in the ciliates^21–23^.

Phagocytosis is ancient, probably having evolved on the order of a billion years ago^24^, and may be the driving force behind eukaryogenesis and the evolution of the nucleus, cytoskeleton, endomembranes, and mitochondrion^25–29^. This view is contested, and some evidence suggests that phagocytosis only fully evolved after the Last Eukaryotic Common Ancestor (LECA) was already established^24,30,31^, and some have conjectured that phagocytosis could only have arisen after the mitochondrion internalized the electron transport chain^32^. The potential for phagocytosis may exist in Asgard archaea^33,34^, but so far hasn’t been observed^35^. The only known case of phagocytosis outside of the eukaryotes is in the planctomycete bacterium Candidatus *Uabimicrobium amorphum*^36^, presumably having evolved convergently.

Supported by prey capture and internal digestion^22^, the core activity of phagocytosis is engulfment of prey. The mechanism of engulfment has been studied extensively in immune cells, with many of the protein players now identified^1,2,37^. Actin-supported membrane extensions (the phagocytic cup) grow around a prey cell, and subsequent membrane shearing and fusion at the phagocytic-cup apex restore a flush plasma membrane and create a topologically separate food vacuole that houses the prey cell.

Despite great progress in the molecular, cell, and evolutionary biology of phagocytosis, the quantitative fitness benefit that phagocytosis bestows upon its host has not been examined. In scenarios of the evolution of phagocytosis, its benefit is simply assumed^25–29^. That phagocytosis, in its current form, provides cells with an advantage seems indisputable given its wide adoption among eukaryotes. How large this advantage is, and whether less effective intermediate forms of phagocytosis are still advantageous, is an open question. Equally unclear is the degree to which the advantage depends on the nature of the ancestor, the environment, and presence of other cellular features such as endomembranes or mitochondria.

Quantifying the *net* fitness contribution of a cellular trait requires the examination of both gross benefit and cost of introducing this new trait. A convenient way of accounting for (some of) the negative effects on fitness is the relative energetic cost of the newly introduced trait, focusing on opportunity costs^38–40^. Such cost-benefit analyses have been carried out for flagella^41,42^, vacuoles^43^, mitochondria^44^, and buoyancy^45^. Here we present a model for quantitating the fitness contribution of phagocytosis, built upon earlier work on pinocytosis and proto-endoplasmic reticulum^46^, that encompasses prey capture, engulfment, and internal and external digestion. The model compares a derived, phagocytosing, state to an ancestral state in which prey is also captured but digested externally.

## Methods

### Outline of the phagocytosis model

We consider a predator cell—the phagocytoser—that captures prey cells, engulfs them, and digests them internally. The fitness of this phagocytoser is compared to an evolutionary precursor cell—the ancestor—that also captures prey but digests these externally.

Both ancestor and phagocytoser are spherical cells of 1000 µm^3^ in volume (typical for eukaryotes^44^) and are limited in their growth rate by nutrient flux. In the ancestor (Figure 1A), prey cells (1 µm^3^ in volume, typical for bacteria^44^) are captured and positioned on the cell surface. Enzymes are excreted by Sec translocases^47^ to convert prey biomass into nutrient molecules that are internalized by nutrient transporters. Both enzymes and nutrients can be lost to the environment.

**Figure 1:**
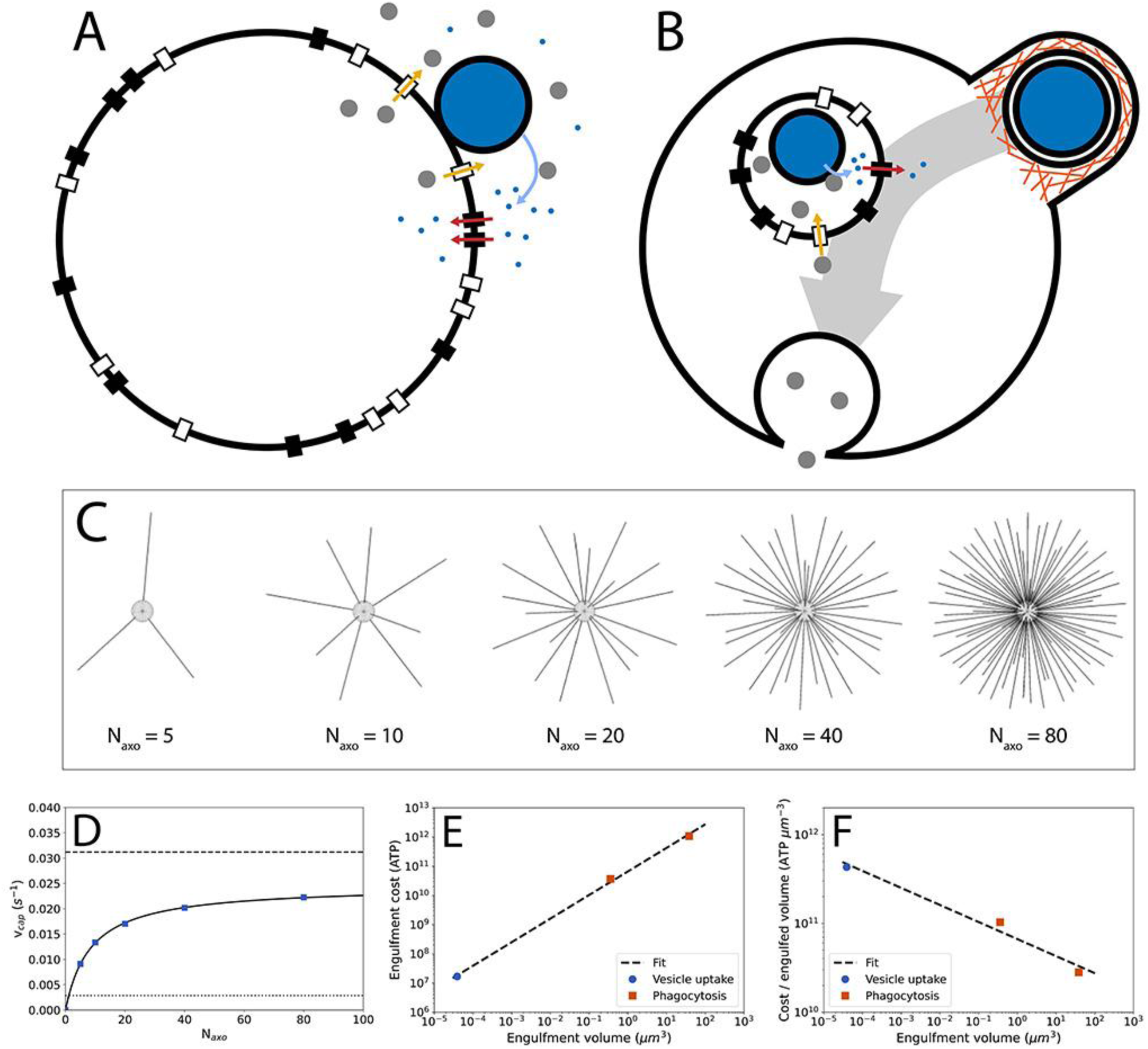
Phagocytosis model, prey capture, and prey engulfment. A) Schematic of the ancestor in the phagocytosis model. Prey capture by axopods is not shown and occurs before the prey cell reaches the surface of the main cell body of the predator. For visual simplicity, only Sec translocases and nutrient transporters close to the prey are shown as active (in the calculations all are active). Large white circle: predator cell; large blue circle: prey cell; grey circles: enzymes; small blue circles: amino acids; black rectangles: amino acid transporters; white rectangles: Sec translocases. B) Schematic of the phagocytoser in the phagocytosis model. Orange lines: actin filaments; small (internal) white circle: food vacuole. C) Smoldyn model of heliozoan prey capture with varying axopod number, *N_axo_*. All axopods have equal lengths. D) Prey capture rate, 𝑣*_cap_*, as a function of axopod number. E) Energy cost of engulfment for phagocytosis and vesicle uptake. Regression: *y* = 6.65 × 10^10^*x*^0.81^. F) Energy cost per engulfed volume for phagocytosis and vesicle uptake. Regression: *y* = 6.65 × 10^10^*x*^−0.19^.

The phagocytoser (Figure 1B) invests in engulfing the prey and digesting it internally to combat the loss of enzymes and nutrients. Prey are captured and subsequently engulfed by a phagocytic cup and internalized in a food vacuole. Sec translocases on the food vacuole, import enzymes into the food vacuole lumen and the nutrients produced by the enzymes are exported into the cytoplasm by nutrient transporters. Finally, the food vacuole fuses with the predator plasma membrane.

### Fitness calculation

Fitness is approximated by two methods (Supplementary Information) that abstract away the details of cellular functioning to serve as a framework for future, more detailed, examinations of the contribution of phagocytosis to fitness. For both methods, we examine growth rate, which is only one aspect of fitness. The first method is a simple ratio of cell division times^46^, in which we assume that other aspects of fitness, i.e. survival, are equal between phagocytoser and ancestor:

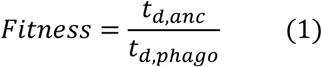

Where *t_d,anc_* and *t_d,phago_* are the cell division times of the ancestor and phagocytoser, respectively. Equation 1 can be applied when the cell surface area is not limiting for growth and survival. In large cells, which have small area-to-volume ratios, the surface area of the cell is a scarce resource and occupying it will come with a steep fitness penalty. To account for this effect (previously included in the cell division time^46^), a second method to calculate fitness is used:

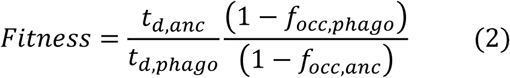

Here, *f_occ,phago_* is the fractional occupancy of the plasma membrane by the phagocytosis and prey capture machinery in the phagocytoser and *f_acc,anc_* is the equivalent for the ancestor. The surface of the cell is essential for its growth and survival, so a fractional decrease of available surface area is likely to have a proportional fractional effect on the fitness. Other relationships between surface area occupancy and the impact of fitness are conceivable and will have to be examined in the future. Unlike the first method, including surface area occupancy can include the fitness contribution of survival, for example if proteins important for survival (e.g. for osmoprotection or toxin expulsion) are displaced from the surface area.

Under the assumption of nutrient limited growth, the cell division time is the ratio of the nutrient requirement for producing a new cell and the nutrient acquisition rate^46^:

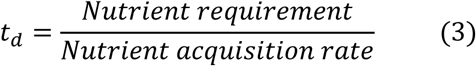

For the ancestor, this becomes (see Supplementary Information):

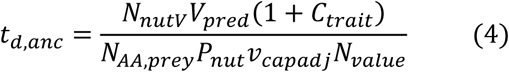

*N_nutV_* is the nutrient requirement per cubic micron of predator volume, *V_pred_* is the predator volume, *C_trait_* is the relative energetic cost of prey capture and external digestion, *N_AAprey_* is the number of nutrients (amino acids) present in a prey cell, *P_nut_* is the probability that a nutrient produced by external digestion reaches the predator, 𝑣*_capadj_* is the prey capture rate adjusted for cell growth (see Supplementary Information), and *N_value_* is the average nutrient value that accounts for a delay in nutrient return to the predator (see Supplementary Information). As an example of loss of nutrient value, consider a batch of nutrient molecules that takes a whole cell division period to return to the predator. It will lose half its value as it will need to supply two cells instead of one.

For the phagocytoser the cell division time becomes:

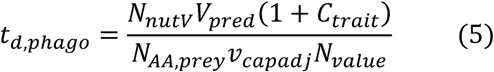

This does not include *P_nut_* because the prey is digested internally and no nutrients are lost. 𝐶_𝑡𝑟𝑎𝑖𝑡_ includes prey capture, engulfment, and internal digestion costs.

### Prey capture

For prey capture, the heliozoan mechanism was used as it allows for the calculation of the relation between energetic cost, cell surface occupancy, and prey capture rate, which are all important for the fitness calculation. For other prey capture mechanisms, such as in ciliates and amoebae, these relationships require more sophisticated models not available at present.

Heliozoan prey capture and subsequent phagocystosis had been observed for *Actinophrys sol* by Albert von Koelliker^4^ in 1848. It involves long cellular protrusions called axopods that capture prey passively and transport it to the predator cell surface to be phagocytosed. Axopods consist of a lipid membrane that is extended out from the plasma membrane by a bundle of microtubules. Axopods can capture a variety of prey of different sizes, including bacteria^48^. Axopod properties, used to calculate energy cost and area occupancy, were extracted from the literature (Figure S1A and S1B, and Table S1 and S2).

Prey capture in the model is based on prey cell diffusion and was quantitatively examined by simulations in the software Smoldyn^49^ (see Supplementary Information, Figure 1C), yielding the relation between the number of axopods, *N_axo_*, and the prey capture rate, 𝑣*_cap_* (Figure 1D):

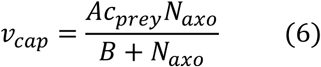

Here, 𝐴 and 𝐵 are fit parameters and 𝑐*_prey_* is the prey concentration. The number of axopods can be related to the axopod energy cost and axopod area occupancy (see Supplementary Information). As the prey concentration is not varied in this study, we haven’t examined saturation of prey uptake rate with prey concentration^50^, and it is assumed that prey concentration is non saturating.

### Prey external digestion

In the ancestor, prey is digested externally by enzymes secreted, at a fixed rate^51^, from plasma membrane-localized protein exporters, called Sec translocases^47^, all over the predator cell surface. The probability with which these enzymes find the prey by random diffusion, *P_enz_*, is determined by Smoldyn simulations^49^ as a function of the prey number that is present on the predator surface simultaneously (Supplementary Information). When only one prey cell is present *P_enz_* = 0.067 (Figure S1C), but this can go up to 1 for larger prey numbers (Figure S1E). Enzymes are assumed to have anchors keeping them attached to the prey, similar to anchoring of enzymes to the predator cell surface^28^. At the prey, the enzymes produce nutrients (amino acids) that reach the predator with a probability, *P_nut_*, of 0.88 (determined by Smoldyn simulation, Figure S1D, Supplementary Information). For simplicity, the prey is assumed to consist entirely of protein, with the rate of amino acid production estimated from digestion rates found in the BRENDA database^52^ (Figure S1F and Table S3). The energy cost of external digestion depends on the number of excreted enzymes, and the area occupancy depends on the number of Sec translocases and nutrient transporters (Supplementary Information). To investigate the limits under which the ancestor can survive, the base numbers for the enzyme to prey probability (*P_enz_*) and the nutrient return probability (*P_nut_*), as calculated above, are multiplied by a factor, 𝑓_𝑚𝑜𝑑_. The factor 𝑓_𝑚𝑜𝑑_ is introduced to adjust *P_enz_* and *P_nut_* simultaneously, as both may depend similarly on fluid flow, confinement, and prey-predator distance. This can’t be accomplished by putting *P_enz_* equal to *P_nut_* because they have different base values, and because *P_enz_* depends on the number of prey cells being digested simultaneously and *P_nut_* doesn’t.

### Prey engulfment

To facilitate estimating the size of the engulfment machinery, and thereby its energetic cost and area occupancy, only protrusion-based engulfment, in which a phagocytic cup emerges from the surface of the predator cell^53,54^, is examined here (Supplementary Information). The energetic cost of engulfment is the sum of the membrane cost and the phagocytic cup plasm cost, with the plasm cost residing in actin filaments^55–59^ and actin accessory proteins (myosin, arp2/3, etc.)^60–62^. In the absence of a quantitative description of the actin cytoskeleton of the phagocytic cup, data were acquired from other membrane structures shaped by the actin cytoskeleton: filopodia^55^, lamellipodia^56^, and endocytic vesicles^57^. This information is combined with energetic cost of membranes^42,63^ and with the geometry and size of phagocytic cups from mouse macrophage^53^ and the choanoflagellate *Codosiga*^54^. In addition to phagocytosis, the energy cost of an endocytic vesicle from *Schizosaccharomyces pombe* ^46,62^ was included in the analysis. The energetic costs calculated here contain only construction costs, operating costs are not accounted for and are expected to contribute only to a minor degree (but this needs to be examined in the future). The energy cost of engulfment is shown in Figure 1E. The cost per unit of engulfed volume (Figure 1F) reveals that larger cups are cheaper per unit of internalized volume, potentially making them easier to evolve than smaller cups and explaining why phagocytic cups are largely absent from small cells.

### Prey internal digestion

The product of engulfment is a food vacuole, consisting of a food vacuole membrane surrounding a prey cell. The food vacuole membrane houses Sec translocases, to import enzymes into the food vacuole lumen, and nutrient transporters, to transport amino acids from the vacuole lumen into the cytoplasm. After the prey is digested, the food vacuole membrane fuses back to the predator plasma membrane and the enzymes, present in the food vacuole lumen, are lost to the external medium. Internal digestion is treated similarly to external digestion except that all the nutrients are recovered and all enzymes make it to the prey (Supplementary Information).

## Results

The cellular traits that define the ancestor and the phagocytoser, and their differences, come with a cost that is accounted for in the 𝐶_𝑡𝑟𝑎𝑖𝑡_ parameter (Eq. 4 and 5). Differences in prey processing machinery lead to differences in the total number of nutrients that need to be accumulated to complete the cell cycle. The nutrient acquisition rate is also different due to the loss of nutrient to the environment by the ancestor, *P_nut_* in equation 4, and due to potential differences in the average nutrient value, *N*_𝑣𝑎𝑙𝑢𝑒_ (related to differences in arrival times of the nutrients, long arrival times lower the nutrient value). Parameter values used to obtain the results are listed in Table S4.

### The limited utility of external digestion

There may be environments in which the phagocytoser can exist, but the ancestor can’t. For example, in the presence of a fluid flow that washes away the enzymes excreted by the ancestor. Similar issues are associated with lifestyles, like swimming, which are more likely to be effective for the phagocytoser than the ancestor. To examine the feasibility of the ancestor, i.e. its ability to grow and reproduce in the absence of the phagocytoser, it is helpful to rewrite the ancestor cell division time from Eq. 4 as (Supplementary Information, Eq. S45, S46, and S47):

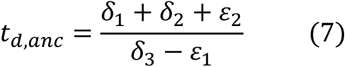

Ancestor cell division time depends on quantities representing the base nutrient requirement (𝛿_1_), the prey capture cost (𝛿_2_), nutrient value discounting (𝜀_2_), the rate of nutrient uptake (𝛿_3_), and the enzyme cost (𝜀_1_). For 𝛿_1_, 𝛿_2_, and 𝜀_2_ the units are nutrients (amino acids) and for 𝛿_3_ and 𝜀_1_ the units are nutrients per second. From equation 7, an immediate insight can be obtained into the feasibility of the ancestor. Enzymes need to be excreted to obtain nutrients, but each enzyme also costs nutrients to produce. The difference 𝛿_3_ − 𝜀_1_ expresses the net nutrient uptake rate, i.e. the rate of nutrient uptake minus the “loss” of nutrients by producing enzymes. If 𝛿_3_ − 𝜀_1_ < 0, an ancestor cell can’t grow, as it would lose more resources in trying to obtain nutrients than it gets back in return. The conditions under which this is true are examined in Fig. 2A, where 𝛿_3_ − 𝜀_1_ is plotted as a function of the time it takes the predator to digest a prey cell, 𝑡_𝑑𝑖𝑔_, and the enzyme and nutrient probability multiplier, 𝑓_𝑚𝑜𝑑_. A lower 𝑓_𝑚𝑜𝑑_ means that the ancestor loses more enzymes and nutrients to the environment. Crucially, 𝛿_3_ depends linearly on 𝑓_𝑚𝑜𝑑_*P_nut_*, and 𝜀_1_ depends inversely on 𝑓_𝑚𝑜𝑑_*P_enz_*. Figure 2A reveals that losses of nutrients and enzymes of approximately 75 to 99% can be tolerated depending on the digestion time. Confined spaces or situations in which kin are in proximity, so that enzymes and nutrients can be shared, increase 𝑓_𝑚𝑜𝑑_ and could allow the ancestor to persist. A longer digestion time, 𝑡_𝑑𝑖𝑔_, improves the prospects of the ancestor because each individual enzyme has more time to release amino acids from the prey, reducing enzyme requirement (and loss) and making digestion more cost efficient.

**Figure 2:**
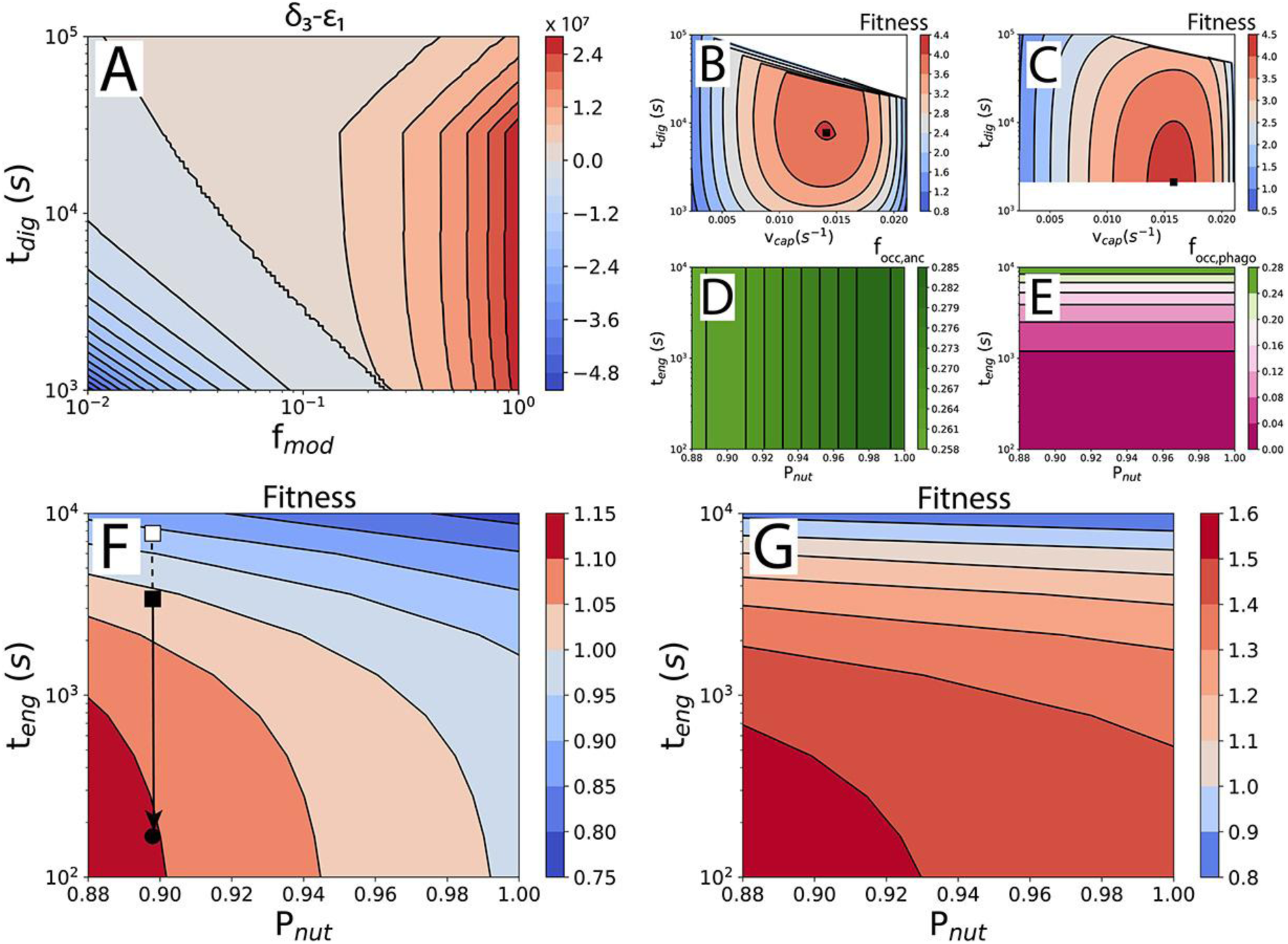
Phagocytosis improves fitness. A) Feasibility of the ancestor. Plot of the net nutrient uptake rate, 𝛿_3_ − 𝜀_1_, as a function of the digestion time, 𝑡_𝑑𝑖𝑔_, and the probability modifier, 𝑓_𝑚𝑜𝑑_ . *P_nut_* is 0.88 and *P_enz_* is 0.21-1.00, depending on 𝑡_𝑑𝑖𝑔_. B) Fitness of ancestral states with different digestion times, 𝑡_𝑑𝑖𝑔_, and prey capture rates, 𝑣*_cap_*, compared to one arbitrary ancestral state from the same set. Result for 𝑡_𝑒𝑛𝑔_ = 167 s and *P_nut_* = 0.88. Black square: maximal fitness, used for the comparison between the ancestor and the phagocytoser in G. Fitness calculated from Eq. 2. White space covers datapoints removed by filters for food vacuole area occupancy, etc. (Supplementary Information). C) Fitness of the phagocytosing states. As in B. D) Fractional occupancy of the cell surface by external digestion and prey capture in the ancestor. E) Fractional occupancy of the cell surface by engulfment and prey capture in the phagocytoser. F) Fitness of the phagocytoser compared with the ancestor, as a function of the nutrient return probability, *P_nut_*, and the engulfment time, 𝑡_𝑒𝑛𝑔_. Calculated from cell division times only (Eq. 1). G) As in F but with fitness calculated from cell division times and surface area occupancy (Eq. 2).

### Fitness of the phagocytoser

The fitness of the phagocytoser can be compared to that of the ancestor if they can both persist in the same niche. Because of the large evolutionary difference between the ancestor and phagocytoser, the fitness must be calculated in two steps. The rationale for the two-step fitness calculation is a separation of the evolutionary timescale over which the parameter values are optimized. Changing the digestion time (𝑡_𝑑𝑖𝑔_) and the prey capture rate (𝑣*_cap_*) involves simple, and therefore rapid, regulatory changes in the number of Sec translocases and axopods. The engulfment time (𝑡_𝑒𝑛𝑔_), on the other hand, is decreased during the process of phagocytosis evolution in which many new proteins are recruited and optimized and therefore takes a long time. The nutrient return probability (*P_nut_*) relates to the nature of the ancestor (and the situation in which it lives) and is therefore not an optimizable parameter.

The first step in the fitness calculation is a comparison of ancestor-to-ancestor, and phagocytoser-to-phagocytoser, in which the parameter values of 𝑡_𝑑𝑖𝑔_ and 𝑣*_cap_* are varied (Figure 2B and 2C), and 𝑡_𝑒𝑛𝑔_ and *P_nut_* are kept constant. One arbitrary combination of 𝑡_𝑑𝑖𝑔_ and 𝑣*_cap_* is chosen as a reference point to which all others are compared. The digestion time optimum found for the phagocytoser, 35 min, is consistent with values reported for ciliates (20 min to 1 h)^22^.

In the second step, the parameter values associated with the fitness optima from the first step (Figure 2B and 2C, black squares) are used to calculate the fitness of the phagocytoser relative to the ancestor. This two-step fitness calculation is repeated for a range of engulfment times, 𝑡_𝑒𝑛𝑔_, and nutrient-return probabilities, *P_nut_* (Figure 2F and 2G).

The phagocytoser beats out the ancestor for shorter engulfment times, 𝑡_𝑒𝑛𝑔_, and lower nutrient return probabilities, *P_nut_* (Figure 2F and 2G). This is true when the fitness is calculated from cell division times alone and even more so when the fitness includes surface area occupancy of the predator plasma membrane. An analysis of which costs subtract most from the fitness is available in Figure S2 and the cell division times are shown in Figure S3. The larger advantage for phagocytosis after accounting for surface area occupancy is caused by the low area occupancy of phagocytosis, an advantage that only starts to disappear at very long engulfment times (Figure 2D and 2E). Area occupancy in the plasma membrane is an important factor in determining fitness when cell surface area comes at a premium, a likely situation for large cells because of small surface-to-volume ratios. This small surface area occupancy of phagocytosis may also explain how single-celled eukaryotes such as ciliates can be almost completely covered in cilia but still phagocytose (Figure S4).

### Fitness of intermediate forms of the phagocytoser

Understanding the evolution of complex traits often encounters the difficulty that all the component parts of the trait have the appearance of being essential, leaving us without an intelligible evolutionary pathway toward the trait. So too for phagocytosis where the production of a food vacuole and its subsequent fusion are both complex processes that appear to be essential for phagocytosis to be effective. It is therefore important to examine the fitness of intermediate states of phagocytosis in which some components of the mechanism are missing or not fully refined. We examine three intermediates: lack of food vacuole fusion, low selectivity for food vacuole membrane proteins, and less effective prey engulfment.

For phagocytosis in which food vacuoles don’t fuse with the plasma membrane (Figure 3A), the food vacuole abundance will reach a steady state, and the cell division time is calculated from (Supplementary Information):

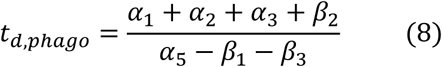

**Figure 3:**
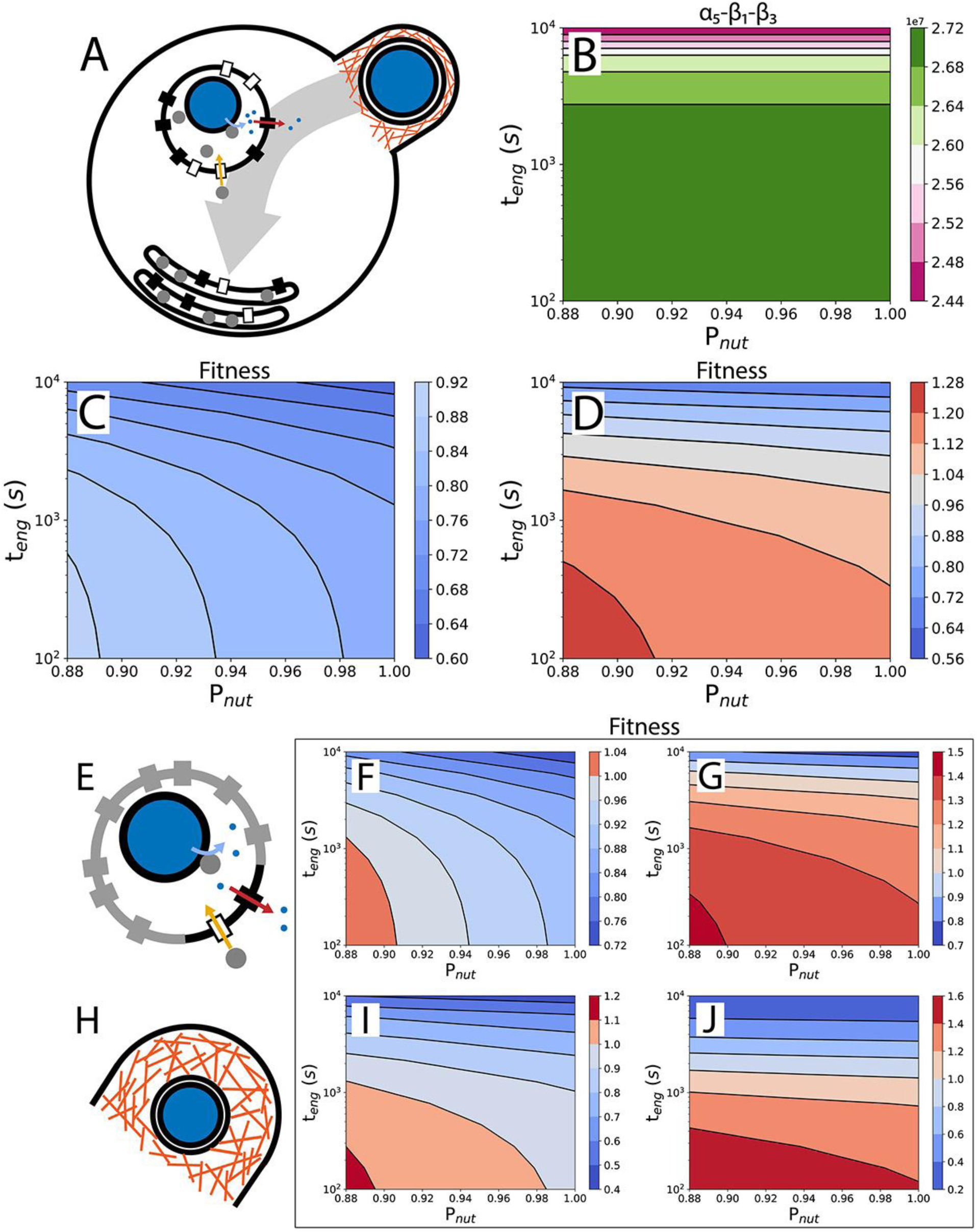
Intermediate forms of phagocytosis improve fitness. A) Schematic of a phagocytoser without food vacuole fusion. Food vacuoles accumulate to a steady state number as flattened membrane compartments. B) Feasibility of phagocytosis without food vacuole fusion. Plot of the net nutrient uptake rate, 𝛼_5_ − 𝛽_1_ − 𝛽_3_, as a function of the engulfment time, 𝑡_𝑒𝑛𝑔_, and nutrient return probability, *P_nut_*. C) Fitness of the phagocytoser compared with the ancestor in the absence of food vacuole fusion, calculated from cell division times only (Eq. 1). D) As in C but with fitness calculated from cell division times and surface area occupancy (Eq. 2). E-G) Fitness calculation in which 90% of the food vacuole surface is occupied by other membrane proteins. E) Schematic of a food vacuole, with membrane proteins other than Sec translocases and nutrient transporters in grey. F) Fitness calculated using cell division times only (Eq. 1). G) Fitness calculated using both cell division time and surface area occupation (Eq. 2). H-J) Fitness calculation for a phagocytoser with larger phagocytic cups. H) Schematic of a larger phagocytic cup. I-J) As in F and G.

The numerator gives the nutrient requirement, which depends on quantities representing the base nutrient requirement (𝛼_1_), the engulfment cost (𝛼_2_), the prey capture cost (𝛼_3_), and nutrient value discounting (𝛽_2_), all expressed in number of nutrients (amino acids). The net nutrient uptake rate, given by the denominator (𝛼_5_ − 𝛽_1_ − 𝛽_3_), must be positive for the phagocytosing organism to grow. This means that the rate of nutrient uptake, 𝛼_5_, must be larger than the sum of the rate of enzyme production (enzymes are lost when the food vacuole membrane fuses with the plasma membrane), 𝛽_1_, and the rate of food vacuole membrane production, 𝛽_3_ (with all three expressed in nutrient number per second). Strikingly, a quantitative examination of the denominator reveals that food vacuole fusion is not required for phagocytosis to persist in the absence of the ancestor because the denominator is always positive (Figure 3B). A phagocytoser without food vacuole fusion can also outperform the ancestor if fractional surface occupancy is important for fitness (Figure 3C & 3D). Volume occupancy of food vacuoles in the cytoplasm is less than 8%, if food vacuoles collapse into a flat shape (Figure 3A).

In the earlier stages of the evolution of phagocytosis, the food vacuole may not have developed a high level of selectivity for the Sec translocases and nutrient transporters required for internal digestion. Consequently, a large part of the food vacuole surface area may be occupied by other proteins (Figure 3E), increasing the digestion time and lowering fitness (Figure 2C). A similar effect may result from a high requirement for receptors to bind the prey cell and recruit the actin cytoskeleton in the formation of the phagocytic cup. Figure 3F & 3G reveal that even with only 10% of the food vacuole membrane available for Sec translocases and nutrient transporters, the phagocytoser can outperform the ancestor.

Prey engulfment will likely be less effective early in the evolution of phagocytosis, which will present itself in longer engulfment time and larger, more energy expensive, phagocytic cups. Fitness quantification can reveal whether a proposed mechanism of engulfment is sufficient to outperform the ancestor (Figure 2F, black square) or not (Figure 2F, white square), depending on the engulfment time that is associated with that mechanism. With subsequent evolution of the engulfment machinery leading to a reduction in the engulfment time and an improvement of the fitness (Figure 2F, arrow and black disc). Modern forms of phagocytosis have engulfment times of minutes^53,64^, similar to that of small-vesicle endocytosis^62^ and prey engulfment by dinoflagellates^65^.

Coordination between actin filaments, a product of the many regulatory proteins associated with the actin cytoskeleton, will initially be lacking, potentially leading to large and expensive phagocytic cups (Figure 3H). Fitness calculations in which the thickness of the phagocytic cup is increased from 0.11 µm to 1 µm reveal that the phagocytoser can still outperform the ancestor but only in a reduced parameter range (Figure 3I & 3J).

## Discussion

### Gradual evolution of phagocytosis

The sequence of cellular transformations by which phagocytosis evolved is unclear, owing in part to its considerable molecular complexity^2,32^. It may have required an intermediate in which prey was partly engulfed^28,66,67^. A sinking-type phagocytosis as seen in excavates^68^ and perhaps present in the last eukaryotic common ancestor^69,70^ may have been an intermediate to the modern diversified forms of phagocytosis across eukaryote diversity^24,31^.

Various studies have revealed how alterations to the mechanism of phagocytosis can affect key parameters identified in our study, such as the engulfment time (𝑡_𝑒𝑛𝑔_, Figure 2F and 2G). Simulations of phagocytosis suggest that it is possible to engulf prey by a receptor only model (i.e. without an actin cytoskeleton), albeit more slowly^37,71^. Knockout of the actin capping protein CapG reduces the uptake rate of zymosan particles by phagocytosis by roughly twofold^72^. Dynamin appears to affect phagocytic cup formation and closure^73,74^. Integrating these results with our phagocytosis model will deepen our understanding of the contribution of the various molecular factors to phagocytosis.

Progress in understanding the evolution of complex traits faces the challenge of the whole trait seemingly having to come into existence all at once with all its components in place. Examples include eyes and wings on the macroscale^75^ and ATP synthase^76^ and flagella^77^ on the molecular and cellular scale. Nevertheless, convincing explanations for the gradual evolution of such complex traits have been proposed^75,77^. A similar difficulty arises in trying to explain the evolution of phagocytosis and it has been claimed that phagocytosis can’t evolve in the absence of the pre-adaptation of vesicle fusion^25,28^. However, as shown here, it is possible for a phagocytoser lacking vesicle fusion to persist (in the absence of the ancestor) and outperform the ancestor, obviating the need for a vesicle fusion pre-adaptation and rendering a gradual transition more plausible.

### Phagocytosis and the endomembrane system

In modern eukaryotes, phagocytosis and other endomembranes always co-occur. The only prokaryote known to perform phagocytosis is a planctomycete that also contains internal membranes^36^. This co-occurrence could mean that phagocytosis is not feasible in the absence of an endomembrane system. The phagocytosis model presented here does not explicitly include other components of the endomembrane system and suggests that phagocytosis could evolve in the absence of such components.

In light of the phagocytosis model, further additions to the endomembrane system become intelligible. For instance, enzymes could be recycled from old to new food vacuoles by a vesicle shunt. This would simultaneously accomplish faster digestion of prey (lowering the key parameter 𝑡_𝑑𝑖𝑔_), because more enzymes are available in the food vacuole at the start, and would prevent the loss of enzymes, as they are removed from the food vacuole before it fuses to the plasma membrane. Similarly, vesicles used for recycling can exchange unnecessary membrane components^2^, such as receptors for prey and the actin cytoskeleton, for components useful in digestion like Sec translocases and nutrient transporters.

It is conceivable that some of the parameter values used in the phagocytosis model depend on the existence of the endomembrane system. Internal digestion in modern phagocytosing species depends on the acidification of the food vacuole lumen by the fusion of endosomes and lysosomes^2^, and it is possible that the rates of prey protein digestion used in the model could only be accomplished in the presence of such acidification. Though presumably a food vacuole could also be acidified by V-type ATPases directly^78^.

The production (or delivery) of membrane necessary for the growth of phagocytic cups is another problem in which the endomembrane system may implicitly underlie the phagocytosis model. The rate of membrane delivery may only be possible in the presence of vesicle fusion to the site of phagocytosis^2,79^, especially in the absence of food vacuole fusion.

It has been argued that phagocytosis is the most complex form of endocytosis and therefore should have arisen after the evolution of small-vesicle endocytosis^32^. The difference in the number of components, ∼127 for phagocytosis and ∼78-95 for small vesicles^32^, only weakly supports such a conclusion. Results from energy cost estimates of phagocytic cups point in the opposite direction: larger phagocytic cups are cheaper per unit internalized volume (i.e. gain) and may thus have an easier time evolving (Figure 1F).

Phagocytosis could have been the starting point of the endomembrane system, as has been argued previously^25–29^. The model presented here shows that phagocytosis can exist in niches in which the ancestor can’t and shows a considerable fitness advantage where they can both occur. This is true for a wide range of parameter values and intermediate states, including an intermediate state in which food vacuoles don’t fuse with the plasma membrane. This suggests that phagocytosis is the more likely candidate for the initiation of endomembrane evolution out of the three alternative scenarios for which fitness has been quantified (Figure 4). The other two scenarios being pinocytosis (of small nutrient molecules) and the proto-endoplasmic reticulum^46^. Comparison with other alternative hypotheses of endomembrane evolution^80–85^ should be carried out in the future but will depend on quantitative formulation of those hypotheses, which has not been attempted yet.

**Figure 4:**
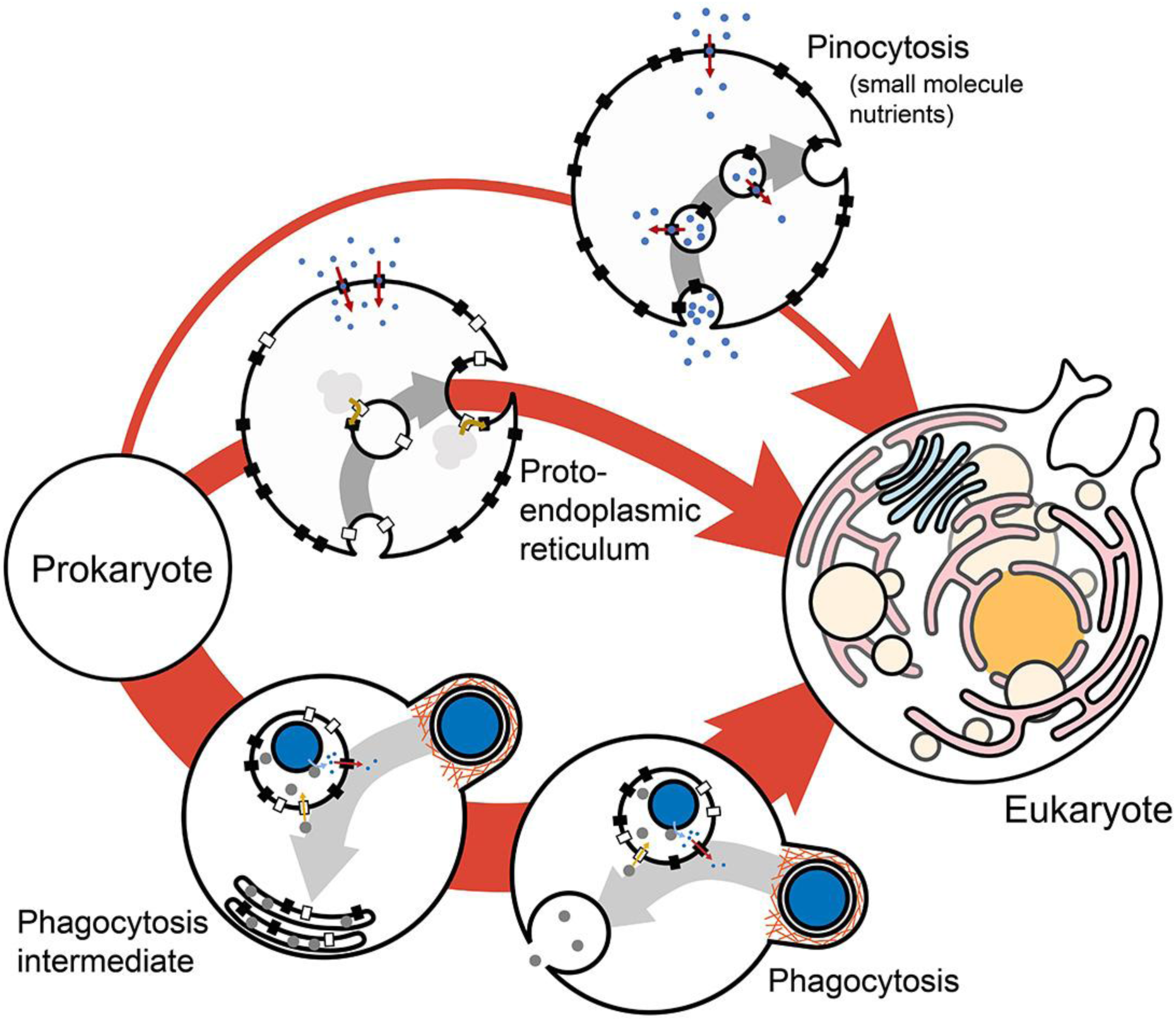
Phagocytosis and the evolution of the endomembrane system. The fitness contribution of phagocytosis can be compared to that of pinocytosis (of small molecules) and proto-endoplasmic reticulum from a previous analysis^46^, showing that phagocytosis is the most likely entry into the evolution of endomembranes and pinocytosis of small molecules the least likely. The likelihood of the evolutionary pathways is proportional to the thickness of the arrows.

### Phagocytosis and mitochondria

Phagocytosis is considered by some to be a precursor to the evolution of mitochondria^27–29,86,87^. Others argue that in the absence of mitochondria, phagocytosis would internalize and digest respiratory enzymes, preventing respiration and phagocytosis from coexisting in the same organism^32^. The logic presented here for food vacuole membrane trafficking and membrane and protein cost doesn’t support respiratory enzyme digestion as they would simply return to the plasma membrane, and even when they don’t, phagocytosis is still beneficial (Figure 3A-3D). It has also been argued that respiration is not required for the existence of phagocytosis as complex phagocytosing organisms exist in anoxic conditions^88,89^. Thus, it seems plausible that phagocytosis could have evolved before mitochondria. Phagocytosis is likely to be more beneficial in large cells and therefore its evolution could have been helped by the presence mitochondria, which would help circumvent an area-to-volume problem in large cells^44^.

## Conclusion

An evolutionary model of phagocytosis is established by which fitness is quantified relative to a non-phagocytic ancestor. This model integrates prey capture, engulfment, and external and internal digestion into a single evolutionary framework, providing a method for judging the biological importance of experimental results that touch on key phagocytosis parameters such as prey engulfment time. It was shown that phagocytosis improves fitness under some conditions and does so even when food vacuoles don’t fuse to the plasma membrane. This work suggests a plausible pathway for the evolution of a complex cellular trait of phagocytosis as well as subsequent evolution of the endomembrane system.

## Acknowledgments

This work was funded by the Moore–Simons Project on the Origin of the Eukaryotic Cell, Simons Foundation 735927, the National Institutes of Health, R35-GM122566, and the National Science Foundation, DBI-2119963.

