## Supplementary Information for "Quantifying the fitness contribution of phagocytosis"

### Calculating fitness of the phagocytosing state

The fitness of a primitive phagocytoser that processes prey internally, is compared with an ancestor that processes prey externally. Building on previous work on the fitness contribution of pinocytosis and proto-endoplasmic reticulum[1] the fitness of a phagocytoser is calculated as:

$$Fitness = \frac{t_{d,anc}}{t_{d,phago}} \qquad (S1)$$

Here, $t_{d,anc}$ is the fitness of the ancestor and $t_{d,phago}$ the fitness of the phagocytoser. In the previous work it was possible to include the fitness effect of area occupancy into the cell division time calculation[1]. Here, the area effect is accounted for heuristically by modifying the ratio of cell division times by a comparison of fractional occupancies of the predator surface area in the phagocytoser, $f_{occ,phago}$, and the ancestor, $f_{occ,anc}$:

$$Fitness = \frac{t_{d,anc}}{t_{d,phago}} \frac{\left(1 - f_{occ,phago}\right)}{\left(1 - f_{occ,anc}\right)} \qquad (S2)$$

The logic behind this modification is that the membrane is an essential bottleneck to the functioning of cells, especially when the predator cells are large, and it is expected that (at least for small fractional occupancies) the negative effect of fitness is linearly proportional to the fractional occupancy. More sophisticated interactions between the fractional occupancies, cell division times, and fitness, are possible but are left to future work.

The cell division times are calculated as a ratio of nutrient requirement and the nutrient acquisition rate, under the assumption that the nutrient acquisition rate is limiting for the growth rate:

$$t_d = \frac{Nutrient\ requirement}{Nutrient\ acquisition\ rate} \qquad (S3)$$

The main driver of predator growth rate is prey capture rate. The difference between the phagocytoser and its externally digesting ancestor is the result of a second-tier effect: the efficiency of processing the prey after it has been captured. This efficiency has three components: (1) loss of nutrients and enzymes, (2) energy cost of processing (excluding enzyme loss), and (3) surface area occupancy of prey processing. The loss of nutrients and enzymes affects nutrient acquisition rate, the energy cost of processing affects the nutrient requirement, and the surface area occupancy (in the present model) affects fitness outside of the cell division time.

### Prey capture (ancestor and phagocytoser)

One of the key parameters that sets the cell division time is the rate of prey capture. In principle we can simply vary the prey capture time and calculate the fitness of the phagocytoser relative to the ancestor purely on differences in prey processing. The reason for including prey capture explicitly is that there is a potential interaction between prey capture and prey processing via the predator cell surface area. A larger area occupancy for prey processing by external digestion compared to by engulfment and internal digestion can affect how much area is available for prey capture. It may also be of interest in the future when comparing to other ancestors that differ in prey capture mechanism or that don't obtain nutrition from prey at all.

Modeling the prey capture rate gives the relation between prey capture rate, energetic cost of the prey capture mechanism, and the area occupancy of the prey capture mechanism. In practice we vary the prey capture rate within a biologically plausible range and calculate the energetic cost and area occupancy form the prey capture rate.

Prey capture mechanisms for single-celled organisms are extremely varied and presumably would have different relationships between capture rate, energetic cost, and area occupancy. The relationship between these quantities can be modeled relatively easily for the heliozoan prey capture mechanism.

#### *Heliozoan prey capture simulation*

Heliozoa are spherical cells covered in radially extending stiff appendages called axopods. Prey cells swim or diffuse into these axopods and get stuck. Subsequently they are transported to the base of the axopods and are phagocytosed. Here, a heliozoan geometry is set up in the reaction-diffusion simulation software Smoldyn[2] and prey cells are produced at a fixed rate per area on a bounding sphere (concentric with the heliozoan cell) with a radius of 200 μm. These prey cells are allowed to diffuse with a diffusion coefficient of 0.396 μm$^2$/s, calculated with the Stokes-Einstein equation:

$$D = \frac{k_B T}{6\pi\eta r_{prey}} \qquad (S4)$$

Using a prey radius $r_{prey}$ of 0.62035 μm (which gives a prey volume of 1 μm$^3$, a temperature T of 298 K, and a viscosity of 8.891x10$^{-4}$ Pa·s.

The number of axopods is varied and the positions of the axopods are assigned automatically using the Fibonacci method giving the coordinates for the base of axopod $i$ as:

$$x_i = \mathrm{r}_{pred}\cos\left(\cos^{-1}\left(2\frac{i}{N} - 1\right) - \frac{\pi}{2}\right)\cos\left(2\pi\frac{i}{\phi}\right) \qquad (S5)$$

$$y_i = \mathrm{r}_{pred}\cos\left(\cos^{-1}\left(2\frac{i}{N} - 1\right) - \frac{\pi}{2}\right)\sin\left(2\pi\frac{i}{\phi}\right) \qquad (S6)$$

$$z_i = \mathrm{r}_{pred}\sin\left(\cos^{-1}\left(2\frac{i}{N} - 1\right) - \frac{\pi}{2}\right) \qquad (S7)$$

Here, $\mathrm{r}_{pred}$ is the predator cell radius (excluding axopods). To obtain the coordinates for the tips of the axopods, the axopod length is added to the predator cell radius. $N$ is the total number of axopods, $i$ is the axopod index and $\phi$ is the golden ratio, $\frac{1+\sqrt{5}}{2}$.

The length of the axopods is 50 μm and the width 0.28 μm. In the simulation the width of the axopods is extended to include the radius of the prey cells and is 0.28 + 2 × 0.62 = 1.52 μm. The reason for this is that the prey cell particles in the Smoldyn simulation don't have a size, only a position. This position is treated as the center point of the prey cell by including the radius of the prey cells in the simulated axopod width. Axopod dimensions are determined from a survey of the literature and are listed in Table S1 and plotted in Figure S1A and S1B.

**Table S1: Cell volumes and axopod dimensions.**

| Species | Cell volume ($\mu m^3$) | Axopod width ($\mu m$)[a] | Axopod length ($\mu m$) | References |
|---|---|---|---|---|
| **Microheliella maris** | 34 | 0.092 | 8 | [3] |
| **Ciliophrys marina** | 1767 | 0.10 | 50 | [4] |
| **Raphidiophrys contractilis** | 3618 | 0.29 | 113 | [5] |
| **Clathrulina elegans** | 9099 | 0.19 | 43 | [6] |
| **Actinophrys sol** | 41441 | 0.49 | 160 | [7] |
| **Actinophrys salsuginosa** | 118847 | 1.64 | 175 | [8] |
| **Actinophrys tauryanini** | 268083 | 4.20 | 292 | [8] |
| **Gymnosphaera albida** | 543480 | 1.25 | 127 | [9] |
| **Actinosphaerium eichhornii** | 8181231 | 0.73 | 148 | [8] |

[a] Axopod width is obtained from electron micrographs where it isn't always clear where along the length of the axopod the image was recorded. As axopods can taper their width accidentally extracting the axopod width from a cross-section closer to the tip could reduce the axopod width estimates.

As soon as a prey cell hits an axopod it is removed from the simulation and counted. Prey cells that diffuse back and hit the bounding sphere are permanently removed from the simulation. This together with constant production of new prey cells at the bounding sphere simulates diffusion through the boundary sphere. The rate at which prey cells are hitting the axopods is the capture rate. Five simulations are performed with different numbers of axopods: 5, 10, 20, 40, and 80. Two additional simulations are performed that don't include axopods. One is a spherical cell in which the surface of the cell is capturing the prey (i.e. a collision of a prey cell with the predator surface counts as a capture). The other is essentially the same but with the surface of the sphere were the tips of the axopods would have been (radius = $r_{pred}$+$length_{axo}$). The simulation is allowed to run for $10^5$ s. The capture rate is obtained over the second half of this period when a stable prey concentration gradient has been established.

### *Capture rate as a function of axopod number*

To incorporate the capture rate results into the phagocytosis model, the number of axopods is plotted against the capture rate and is fitted with:

$$v_{cap} = \frac{A c_{prey} N_{axo}}{B + N_{axo}} \qquad (S8)$$

Here, $A$ is a fit parameter that together with the prey concentration, $c_{prey}$, gives the maximum capture rate and $B$ is the equivalent of the Michaelis constant in the Michaelis-Menten equation. To obtain the value of A from the fit, the prey concentration at infinite distance, $c_\infty$, is required from the simulation. The simulation itself doesn't include prey concentration at infinite distance *per se* but this can be calculated from the flux through the surface of a sphere that is concentric with the cell. This is obtained from the simulation without axopods and with a sphere radius of $r_{pred}$+$length_{axo}$, using the flux through a bounding sphere[10] at distance r:

$$J(r) = -\frac{Dc_\infty a}{r^2} \qquad (S9)$$

D is the diffusion coefficient of the prey cell, $c_\infty$ is the prey concentration at infinite distance, $a$ is the radius of the absorbing boundary, and $r$ is the radius of the sphere at which the flux is examined. Setting r=a and rewriting yields:

$$c_\infty = \frac{J(a)a}{D} \qquad (S10)$$

Here, the minus sign has been omitted because it is only relevant for the direction of the flux. $J(a)$ is the flux (per unit surface area) over the surface of the absorbing sphere (i.e. the number of prey cells hitting the surface per square micron per second). This is obtained from the simulation (taking the absolute value). Once the $c_\infty$ is obtained from the simulations the parameter $A$ can be determined from equation S8 by substituting $c_\infty$ for $c_{prey}$. Once $A$ is found it can be combined with $B$ is any desired value of $c_{prey}$ and $N_{axo}$ to determine the capture rate. In practice, the phagocytosis model varies the capture rate, $v_{cap}$, and the number of axopods, $N_{axo}$, is determined from:

$$N_{axo} = \frac{B}{\frac{Ac_{prey}}{v_{cap}} - 1} \qquad (S11)$$

This can be multiplied with the cost, $C_{axo}$, or surface area occupancy, $A_{axo}$, of a single axopod to determine the total cost and area occupancy of the prey capture mechanism.

#### *Energy cost of prey capture*

The cost of a single axopod includes the membrane surrounding the axopod, the microtubules in the axopod, and the microtubules extending from the middle of the cell to the base of the axopod.

$$C_{axo} = C_{mem} + C_{MT} \qquad (S12)$$

Here, $C_{mem}$ is the cost of the membrane and $C_{MT}$ the cost of the microtubules. The membrane cost can be determined from the per unit area cost of a membrane, $C_{area}$, which includes lipids and proteins[11] and by assuming a cylindrical axopod shape:

$$C_{mem} = L_{axo} 2\pi r_{axo} C_{area} \qquad (S13)$$

Here, $L_{axo}$ is the length of the axopod and $r_{axo}$ is the radius of the axopod. The membrane area at the axopod tip is assumed to be equivalent to the membrane area that is removed from the plasma membrane at the base so that only the sides of a cylinder contribute to the axopod membrane cost.

The cost of the microtubules is calculated from the axopod cross-sectional area and the number of microtubules per unit cross-sectional area (microtubule density). The microtubule density is extracted from electron micrographs (Table S2). The microtubule cost includes the segments that extend from the cell surface to the cell center and is given by:

$$C_{MT} = \frac{N_{MT} L_{MT} N_{pf} N_{AA,tub} C_{AA}}{d_{monomer}} \qquad (S14)$$

Here, $N_{MT}$ is the number of microtubules in an axopod (calculated from the microtubule density and axopod cross-sectional area), $L_{MT}$ is the length of the microtubules (equal to $L_{axo} + r_{predator}$), $N_{pf} = 13$ is the number of protofilaments in a microtubule[11], $N_{AA,tub} = 447$ is the average number of amino acids in a tubulin monomer (averaged over α- and β-tubulin)[11], $C_{AA}$ is the energy cost per average amino acid, and $d_{monomer} = 0.004\ \mu m$ is the distance a single monomer spans along the length of the microtubule[11]. $C_{mem}$ and $C_{MT}$ can be combined to calculate the cost of an individual axopod, $C_{axo}$. Multiplying the cost of an individual axopod by the number of axopods gives the total cost of prey capture (in units of ATP):

$$C_{cap} = C_{axo} N_{axo} \qquad (S15)$$

*Area occupancy of prey capture*

The area occupancy of the axopods on the surface of the predator cell is calculated by combining the cross-sectional area with the axopod number:

$$A_{axo,all} = N_{axo} A_{axo} = \frac{B}{\frac{Ac_{prey}}{v_{cap}} - 1} \pi {r_{axo}}^2 \qquad (S16)$$

With $r_{axo}$ being the axopod radius. Giving the fractional occupancy of prey capture as:

$$f_{occ,cap} = \frac{A_{axo,all}}{A_{pm}} \qquad (S17)$$

$A_{pm}$ is the area of the predator plasma membrane at the start of the cell division period.

**Table S2: Microtubule (MT) density in axopod cross-sections.**

| Species | MT density ($\mu m^{-2}$) | References |
|---|---|---|
| Actinosphaerium nucleofilum | 655 | [12] |
| Raphidiophrys elegans | 796 | [13] |
| Dimorpha mutans | 1006 | [13] |
| Average | **819** | |

### External digestion (ancestor)

In the model, both the ancestor and the phagocytoser have the same heliozoan prey capture mechanism. The difference is in how the captured prey is processed. In the ancestor, the prey is held at the cell surface and enzymes are excreted all over the surface of the predator cell. Some of these enzymes make it to the prey, with probability $P_{enz}$, and the remainder are lost to the environment. The enzymes that make it to the prey stay on the prey until the prey is completely digested. The nutrients produced by the enzymes diffuse to the predator cell with probability $P_{nut}$, with the remainder being lost to the environment.

#### *Calculation of $P_{enz}$ and $P_{nut}$*

$P_{enz}$ and $P_{nut}$ are determined using Smoldyn simulations[2]. For $P_{enz}$ there is a large sphere that represents the predator cell and touching it is a small sphere representing the prey cell (Figure S1C). The axopods are not included in these simulations. A thousand particles representing the enzymes are positioned at random over the entire surface of the predator cell. These are released and diffuse with a diffusion coefficient of 85 μm$^2$/s.[14] As an enzyme particle hits the surface of the prey cell it is removed from the simulation. The number of remaining particles is tracked in time and reaches a plateau (Figure S1C). $P_{enz}$ is calculated by comparing the plateau to the starting enzyme number: $P_{enz} = \frac{N_{start} - N_{plateau}}{N_{start}}$. To determine $P_{nut}$, an equivalent simulation is performed but now the particles start on the prey sphere and diffuse with a diffusion coefficient of 750 μm$^2$/s (Figure S1D). Simulations were performed three times for both $P_{enz}$ and $P_{nut}$ with the average value being used in the phagocytosis model.

When many prey are caught in a short time window and/or prey processing takes a long time, there are multiple prey present at the predator surface at any given moment (Figure S1E). This raises $P_{enz}$ because there is a larger target for the enzymes. The relation between $P_{enz}$ and the number of prey is determined by repeating the simulations as described but with the number of prey cells varying between 1 and 128 (Figure S1E), positioning prey equidistantly on the surface of the predator cell using the Fibonacci method already described for the Heliozoan prey capture mechanism. This simulation data was fitted with:

$$P_{enz} = \frac{A_{penz} N_{prey}}{B_{penz} + N_{prey}} \qquad (S18)$$

Here, $A_{penz} = 1.112$ and $B_{penz} = 13.71$ are fit parameters. This equation is used in the phagocytosis model with the additional rule that if the $P_{enz}$ given by this equation is higher than 1, a value of exactly 1 is used.

#### *Calculating the digestion time, $t_{dig}$*

Enzymes accumulate on the prey, and turn the prey biomass, which is considered to consist of pure protein, into amino acids. This digestion takes a time $t_{dig}$ to complete. The digestion time depends on the number of amino acids to be released from the prey, $N_{AA,prey}$, and the number of enzymes accumulating over time. The $N_{AA,prey}$ is given by the integral:

$$N_{AA,prey} = \int_0^{t_{dig}} N_{enz}(t) P_{enz} k_{prot} dt = \frac{N_{Sec} k_{prot} P_{enz}}{t_{ins}} \int_0^{t_{dig}} t dt \qquad (S19)$$

Here, $N_{enz}(t)$ is the number of enzymes excreted by the predator cell for this particular prey after time $t$, $N_{Sec}$ is the number of Sec translocases devoted to the excretion of enzymes, $k_{prot}$ is the amino acid production rate per enzyme, and $t_{ins}$ is the time it takes for a single Sec translocon to excrete a single enzyme. Solving the integral yields:

$$N_{AA,prey} = \frac{N_{Sec} k_{prot} P_{enz} {t_{dig}}^2}{2 t_{ins}} \qquad (S20)$$

Rewriting gives the digestion time:

$$t_{dig} = \sqrt{\frac{2 t_{ins} N_{AA,prey}}{N_{Sec} k_{prot} P_{enz}}} \qquad (S21)$$

In the phagocytosis model $t_{dig}$ is a variable, but this relation is used to find the number of Sec translocases $N_{Sec}$.

#### *Calculating the energy cost of external digestion*

The enzymes used to degrade prey cells come with an energy cost. The total energy expenditure on enzymes depends on how many enzymes are excreted by the predator. Enzymes are excreted by the predator throughout the digestion time and digestion events happen throughout the cell division period. Each individual enzyme comes with a cost of $N_{AA,enz} C_{AA}$, i.e. the number of amino acids in an enzyme multiplied by the energetic cost of an (average) amino acid. The number of enzymes produced to digest a single prey cell can be obtained from:

$$N_{AA,prey} = \int_0^{t_{dig}} N_{enz}(t) P_{enz} k_{prot} dt = k_{enz} P_{enz} k_{prot} \int_0^{t_{dig}} t dt \qquad (S22)$$

With $k_{enz}$ being the rate of enzyme excretion (considering only a single prey cell). Solving the integral and rewriting yields:

$$k_{enz} = \frac{2 N_{AA,prey}}{P_{enz} k_{prot} {t_{dig}}^2} \qquad (S23)$$

The number of enzymes produced over the whole digestion period, $N_{enz,prey}$, is obtained by multiplying $k_{enz}$ by $t_{dig}$:

$$N_{enz,prey} = \frac{2 N_{AA,prey}}{P_{enz} k_{prot} t_{dig}} \qquad (S24)$$

The cost of digesting a single prey cell is equal to the product of the number of enzymes required to digest a single prey cell, $N_{enz,prey}$, and the cost per enzyme, $N_{AA,enz} C_{AA}$. The total number of prey capture events can be obtained from the prey capture rate and the cell division time, $v_{capadj} t_{d,anc}$, where $v_{capadj}$ is the prey capture rate adjusted for predator growth (see "Correcting $v_{cap}$ for growth" section) and $t_{d,anc}$ is the cell division time of the ancestor. The absolute energetic cost of external digestion, expressed in units of ATP, is given by:

$$C_{dig,ext} = \frac{2N_{AA,prey}}{P_{enz}k_{prot}t_{dig}} N_{AA,enz}C_{AA}v_{capadj}t_{d,anc} \qquad (S25)$$

*Calculating the area occupancy of Sec translocases and nutrient transporters*

The amino acids freed from prey biomass by the enzymes are imported by nutrient transporters. Energetic costs of both Sec translocases and nutrient transporters are not accounted for explicitly because they are embedded in the membrane, displacing what was already there and therefore not contributing additional energetic costs. Although Sec translocases and nutrient transporters don't contribute to the energetic cost of external digestion, they do impact the fitness of the ancestor by occupying the predator plasma membrane.

The total area occupied by Sec translocases and nutrient transporters depends on their copy numbers in the membrane. The number of Sec translocases for a single prey cell is given by:

$$N_{Sec} = k_{enz}t_{ins} = \frac{2N_{AA,prey}t_{ins}}{P_{enz}k_{prot}{t_{dig}}^2} \qquad (S26)$$

The number of nutrient transporters depends on how many nutrients are produced per unit time by the enzymes. The number of enzymes is a function of the number of Sec translocases. The required number of nutrient transporters will vary over time, as enzymes accumulate on the prey, but in the model the nutrient transporter number is constant at the maximum requirement (achieved at $t_{dig}$). The number of amino acids being produced by the enzymes at time $t_{dig}$ is $\frac{N_{Sec}t_{dig}k_{prot}}{t_{ins}}$. Not all amino acids make it back to the cell, giving $\frac{N_{Sec}t_{dig}k_{prot}P_{nut}}{t_{ins}}$ for the number of amino acids making it to the predator. It is assumed that these nutrients are transported by the nutrient transporters at perfect efficiency which gives a transport rate of $N_{trans}k_{trans}$ per prey cell. Where $N_{trans}$ is the number of nutrient transporters per prey cell and $k_{trans}$ is the transport rate per nutrient transporter. Equating nutrient production (including $P_{nut}$) and nutrient transport we obtain:

$$\frac{N_{Sec}t_{dig}k_{prot}P_{nut}}{t_{ins}} = N_{trans}k_{trans} \qquad (S27)$$

This allows us to calculate the number of nutrient transporters, $N_{trans}$, as a function of the number of Sec translocases, $N_{Sec}$:

$$N_{trans} = \frac{N_{Sec}t_{dig}k_{prot}P_{nut}}{t_{ins}k_{trans}} = \frac{2N_{AA,prey}P_{nut}}{P_{enz}t_{dig}k_{trans}} \qquad (S28)$$

The number of prey present at any given moment is given by $t_{dig}v_{cap}$. Here, $v_{cap}$ is used instead of $v_{capadj}$ because the area occupied by Sec translocases and nutrient transporters is compared to the predator plasma membrane area at the start of the cell cycle. The area occupied by a single Sec translocase or nutrient transporter is $A_{mp}$. The lipids surrounding the Sec translocases and Nutrient transporters are included in the required area by using the fraction of membrane area that is membrane protein, $f_{memprot}$. Following these considerations the fractional occupancy of the ancestor plasma membrane by Sec translocases and nutrient transporters can be calculated as:

$$f_{occ,ext} = \frac{1}{A_{pm}} \frac{A_{mp}}{f_{memprot}} (N_{trans} + N_{Sec}) t_{dig} v_{cap} \qquad (S29)$$

$A_{pm}$ is the area of the predator plasma membrane at the start of the cell division period. Finally, the fractional occupancy of ancestor plasma membrane is obtained by including prey capture:

$$f_{occ,anc} = f_{occ,ext} + f_{occ,cap} \qquad (S30)$$

### Engulfment (phagocytoser)

The phagocytoser differs from the ancestor in how the captured prey is processed. After the prey is captured, it is positioned on the surface of the predator plasma membrane where it is engulfed by a phagocytic cup. From the phagocytic cup a food vacuole forms in which the prey is digested (see section on internal digestion). The phagocytic cup comes with an energy cost and an area occupancy.

#### *Energy cost of engulfment*

The phagocytic cup starts a circular protrusion of the predator cell membrane which forms around the prey cell. This grows until it completely envelops the prey cell. The phagocytic cup comes together at the apex of the prey where it fuses to produce a new patch of plasma membrane from its external facing membrane and a food vacuole membrane from the prey-facing membrane. The phagocytic cup is composed of membrane, actin, and actin accessory proteins.

The membrane cost is determined by multiplying the total membrane area by the per unit area energy cost of a membrane, $1.45 \times 10^9$ ATP $\mu m^{-2}$ (including both lipids and membrane proteins)[11].

Actin abundances per unit volume are obtained from structures that are similar to phagocytic cups, i.e. actin-based structures that manipulate membranes. Estimates of actin abundance and structure dimensions were obtained for a filopodium[15], lamellipodium[16], and for endocytic vesicle construction[17]. For the lamellipodium calculation monomer number per unit actin filament length was obtained from[18,19]. From the actin abundance, densities of actin monomers were calculated of $1.33 \times 10^5$ $\mu m^{-3}$ for the filopodium, $5.85 \times 10^5$ $\mu m^{-3}$ for the lamellipodium, and $1.22 \times 10^6$ $\mu m^{-3}$ for the endocytic vesicle. The cost of actin per cubic micron is given by:

$$C_{actin} = d_{actin} N_{AA,actin} C_{AA} \qquad (S31)$$

Here, $d_{actin}$ is the density of actin monomers, $N_{AA,actin}$ is the number of amino acids per actin monomer (376 amino acids), and $C_{AA}$ is the energy cost per average amino acid. This leads to a geometric mean actin energy cost of $4.97 \times 10^9$ ATP $\mu m^{-3}$.

The abundance of actin accessory proteins is estimated from three different sources. The first is the abundance[20] of actin associated proteins compared to actin[21] in whole cells of *Dictyostelium discoideum*. If the elongation factors are excluded from the list of actin associated proteins the abundance of actin + actin associated proteins is 4-fold that of actin alone. The second and third estimates of actin accessory protein cost, which accounts for differences in protein amino acid length, come from endocytic vesicle construction in *Schizosaccharomyces pombe*[22]. The second estimate uses the total cytoplasmic abundance of actin compared to that of other proteins involved in endocytic vesicle construction and results in an energy cost multiplier of 3.4-fold. The third estimate derives from the abundance of actin and accessory proteins at the site of endocytic vesicle construction and results in an energy cost multiplier of 2.4. This gives an average multiplier of 3.3-fold and an energetic cost of actin plus accessory proteins of 1.6x$10^{10}$ ATP $\mu m^{-3}$.

The membrane and phagocytic cup plasm costs (actin plus accessory proteins) can be combined with information about the size and shape of phagocytic cups to obtain an estimate of the energy cost of engulfment. The calculation is performed for two phagocytic cups of differing size and shape, one from the choanoflagellate *Codosiga*[23] and the other from a mouse macrophage[24]. For the choanoflagellate this yields a phagocytic cup cost of 3.7x$10^{10}$ ATP for an enclosed volume (or prey volume) of 0.36 $\mu m^3$, giving an energy cost per unit enclosed volume of 10.0x$10^{10}$ ATP $\mu m^{-3}$. For the mouse macrophage these numbers are 1.1x$10^{12}$ ATP, 38 $\mu m^3$, and 2.8x$10^{10}$ ATP $\mu m^{-3}$.

These estimates are combined with the energy cost of producing an endocytic vesicle[1]. For the endocytic vesicle we have a vesicle construction cost of 1.7x$10^7$ ATP, an enclosed volume of 3.9x$10^{-5}$, and an energy cost per unit enclosed volume of 4.3x$10^{11}$ ATP $\mu m^{-3}$. A regression is performed on these three datapoints yielding the energy cost of engulfing a volume as:

$$C_{eng,single} = 6.65 \times 10^{10} {V_{eng}}^{0.808} \qquad (S32)$$

The engulfment cost, $C_{eng,single}$, has units of ATP and the engulfment volume, $V_{eng}$, has units of $\mu m^3$. To obtain the total cost of engulfment the cost of a single engulfment is multiplied by the number of simultaneous engulfment events, $t_{eng} v_{capadj}$:

$$C_{eng} = C_{eng,single} t_{eng} v_{capadj} \qquad (S33)$$

*Area of engulfment*

The area occupancy of the phagocytic cup is obtained by assuming a spherical prey with a volume of 1 $\mu m^3$ and a phagocytic cup thickness of 0.11 µm obtained from the choanoflagellate[23]. This yields an area occupancy, $A_{phago}$, of 1.67 $\mu m^2$, which is divided by two to account for growth and decline. It is assumed that the outer surface of the phagocytic cup can't be used by any other cellular processes.

The fractional area occupancy for the phagocytoser includes prey capture and engulfment and is given by:

$$f_{occ,phago} = \frac{\frac{1}{2} A_{phago} t_{eng} v_{capadj} + A_{axo,all}}{A_{pm}} \qquad (S34)$$

$A_{phago}$ is the area occupancy of a single phagocytic cup at maximum extent, $t_{eng}$ is the time of engulfment, $v_{capadj}$ is the growth adjusted prey capture rate, $A_{axo,all}$ is the area occupancy of prey capture, and $A_{pm}$ is the plasma membrane area of the predator.

#### *Difference in energetic cost per engulfment volume*

The choanoflagellate phagocytic cup is 3.6-fold as expensive, per engulfment volume, as the mouse macrophage. This is due to a 1.8-fold increase in phagocytic cup plasm cost and a 12.5-fold increase in membrane cost. Given the engulfment volumes involved, the change in membrane cost expected from a difference in surface area to volume alone, assuming spherical prey, is 6.8-fold, with the remaining 1.8-fold being due to differences in phagocytic cup shape. Thus, the choanoflagellate cup is more expensive mostly because of its small size.

### Internal digestion in a food vacuole (phagocytoser)

Upon completion of prey engulfment, a food vacuole is formed in the predator cytoplasm. The food vacuole membrane contains Sec translocases that specifically transport enzymes to convert the prey biomass into amino acids (as with external digestion, it is assumed that prey is composed entirely of protein). These amino acids are transported from the food vacuole lumen and into the cytoplasm by nutrient transporters. The key parameter controlling the internal digestion process is the digestion time, $t_{dig}$. This sets the requirement for Sec translocases and nutrient transporters, it determines the efficiency of digestion (by setting the time that individual enzymes spend on the prey), and it determines the contribution of the food vacuole membrane energy cost to the predator whole cell cost.

#### *Digestion time, $t_{dig}$*

The digestion time is calculated similar to that of the ancestor, except that all enzymes make it to the prey (i.e. $P_{enz} = 1$), yielding:

$$t_{dig} = \sqrt{\frac{2t_{ins}N_{AA,prey}}{N_{Sec}k_{prot}}} \qquad (S35)$$

As with the ancestor, $t_{dig}$ is treated as a variable in the phagocytosis model, but this equation is used to obtain the number of Sec translocases, $N_{Sec}$.

#### *Cost of food vacuole membrane*

The energy cost of the food vacuole membrane is a product of the food vacuole membrane area, a constant related to the prey volume, $V_{prey}$, and the energy cost per unit membrane area, $C_{area}$. To account for the number of food vacuoles being present at any given moment, this is multiplied by $t_{dig}v_{capadj}$, yielding:

$$C_{fv} = 4\pi\left(\frac{3V_{prey}}{4\pi}\right)^{\frac{2}{3}} C_{area}t_{dig}v_{capadj} \qquad (S36)$$

*Cost of enzymes*

The energy cost of the enzymes that digest the prey inside of the food vacuoles needs to be included in the total cost of internal digestion. This differs from external digestion in that no enzymes are lost before they are used, but it is similar in that the enzymes are ultimately lost. Enzymes are lost because in the primitive phagocytosis examined here, there is no mechanism for recycling the enzymes, they are lost to the external medium upon food vacuole fusion with the plasma membrane after the prey has been digested. The total energy cost of the enzymes used for internal digestion can be calculated similarly to that for external digestion except that $P_{enz} = 1$, yielding:

$$C_{enz,int} = \frac{2N_{AA,prey}}{k_{prot}t_{dig}} N_{AA,enz}C_{AA}v_{capadj}t_{d,phago} \qquad (S37)$$

Like for external digestion the total cost depends on the cell division time, $t_{d,phago}$.

### Predator energy budget and correcting for growth

The energetic costs calculated so far are absolute costs of constructing the various cellular traits, expressed in units of ATP. Only by normalizing these costs to the cellular budget can these absolute costs be made relevant for the calculation of fitness. Here, we use the cell budget before any of the cellular traits discussed here (prey capture, engulfment, etc.) are added, $C_{budget}$, to calculate a relative energetic cost. $C_{budget}$ is obtained from the predator volume by a simplified empirical relation[25]:

$$C_{budget} = 2.7 \times 10^{10} V_{pred}{}^{0.97} \qquad (S38)$$

As the cell cycle progresses the predator cell grows as does the cell budget. To account for this an average cell budget can be calculated. The average value depends on the pattern of growth, exponential, linear, etc. Here, the pattern of growth is set by the prey capture mechanism. It can be assumed that the number of axopods doubles over the cell division period, but the rate at which axopods are added depends on how the capture rate changes with the axopod number. At optimal axopod numbers, the doubling of axopod number during cell division changes the prey capture rate only marginally. For this reason and for the sake of simplicity the growth is assumed to be linear. This means that the average cellular energy budget, given by $C_{budget}f_{budget}$, can be calculated by setting $f_{budget} = 1.5$.

### Correcting $v_{cap}$ for growth

As the predator grows, its capacity to capture prey, and accumulate biomass, increases with the rising number of axopods. This effect can be incorporated into the phagocytosis model by assuming a linear rate of increase in the number of axopods and then taking the average over the prey capture rate:

$$v_{capadj} = \frac{1}{N_{axo}{}^*} \int_{N_{axo}{}^*}^{2N_{axo}{}^*} \frac{Ac_{prey}N_{axo}}{B + N_{axo}} dN_{axo} \qquad (S39)$$

${N_{axo}}^*$ is the number of axopods at the start of the cell cycle and is given by:

$$ {N_{axo}}^* = \frac{v_{cap}B}{Ac_{prey} - v_{cap}} \qquad (S40) $$

Here, $c_{prey}$ is the prey concentration far from the predator cell. Solving the integral yields:

$$ v_{capadj} = \frac{Ac_{prey}}{{N_{axo}}^*}\left({N_{axo}}^* + Bln({N_{axo}}^* + B) - Bln(2{N_{axo}}^* + B)\right) \qquad (S41) $$

This is used to calculate $v_{capadj}$ from $v_{cap}$.

### Delayed nutrient return penalty

For both the ancestor and the phagocytoser the digestion time is varied. The longer the digestion time the more efficient the production of amino acids by enzymes, i.e. fewer enzymes convert the same amount of biomass into amino acids but over a longer period. However, if an amino acid is produced and returned to the cell late, this reduces its value because the cell has increased in volume in the meantime. Thus, a larger $t_{dig}$ also reduces the value of an amino acid. This effect is accounted for by a delayed return penalty. The degree to which the nutrient value is reduced depends on $t_{dig}$ and the cell division time. Assuming a linear increase in volume over the cell division time the nutrient value is given by:

$$ Nutrient\ value = 1 - \frac{1}{2}\frac{t_{dig} - t_\beta - t_\alpha}{t_d} \qquad (S42) $$

Here, $t_\beta$ is the time since the start of digestion that a particular enzyme has been produced by the predator (diffusion time of enzyme to prey is fast and therefore ignored), and $t_\alpha$is the time since the moment that the enzyme was produced that a nutrient, produced by that enzyme, arrives at the predator cell. The delayed return penalty, or average nutrient value, $N_{value}$, is calculated by integrating over all enzymes and all nutrients:

$$ N_{value} = \frac{\int_0^{t_{dig}}\int_0^{t_{dig}-t_\beta}\left(1 - \frac{1}{2}\frac{t_{dig} - t_\beta - t_\alpha}{t_d}\right)dt_\alpha dt_\beta}{\int_0^{t_{dig}}\int_0^{t_{dig}-t_\beta} 1\, dt_\alpha dt_\beta} \qquad (S43) $$

Here, $\int_0^{t_{dig}}\int_0^{t_{dig}-t_\beta} 1\, dt_\alpha dt_\beta$ is a normalization constant. Solving the integrals yields:

$$ N_{value} = 1 - \frac{t_{dig}}{6t_d} \qquad (S44) $$

### Calculation of $t_{d,anc}$ and $t_{d,phago}$

The cell division time of the ancestor can be calculated by adding all the costs together to get a nutrient requirement and modifying the nutrient uptake rate by the delayed nutrient return penalty. This yields an ancestral division time of:

$$t_{d,anc} = \frac{N_{nutV}V_{pred}(1 + C_{trait})}{N_{AA,prey}P_{nut}v_{capadj}N_{value}} \qquad (S45)$$

$N_{nutV}$ is the nutrient (or amino acid) requirement per unit predator volume, $V_{pred}$ is the predator volume, $C_{trait}$ is the cost of prey capture and external digestion, and $N_{AA,prey}P_{nut}v_{capadj}$ is the number of nutrient molecules arriving per unit time. Substituting the trait costs and $N_{value}$ yields:

$$t_{d,anc} = \frac{\delta_1 + \delta_2 + \varepsilon_1 t_{d,anc}}{\delta_3\left(1 - \frac{t_{dig}}{6t_{d,anc}}\right)} = \frac{\delta_1 + \delta_2 + \varepsilon_1 t_{d,anc}}{\delta_3 - \frac{\delta_3 t_{dig}}{6}\frac{1}{t_{d,anc}}} = \frac{\delta_1 + \delta_2 + \varepsilon_1 t_{d,anc}}{\delta_3 - \frac{\varepsilon_2}{t_{d,anc}}} \qquad (S46)$$

$$\delta_1 = N_{nutV}V_{pred} \qquad (S46a)$$

$$\delta_2 = N_{nutV}V_{pred}C_{cap}\frac{1}{C_{budget}} \qquad (S46b)$$

$$\delta_3 = N_{AA,prey}P_{nut}v_{capadj} \qquad (S46c)$$

$$\varepsilon_1 = N_{nutV}V_{pred}\left(\frac{2N_{AA,prey}}{P_{enz}k_{prot}t_{dig}}N_{AA,enz}C_{AA}v_{capadj}\right)\frac{1}{C_{budget}f_{budget}} \qquad (S46d)$$

$$\varepsilon_2 = N_{AA,prey}P_{nut}v_{capadj}\frac{t_{dig}}{6} \qquad (S46e)$$

The ancestral cell division time can be obtained from the deltas and epsilons by rewriting the equation:

$$t_{d,anc} = \frac{\delta_1 + \delta_2 + \varepsilon_2}{\delta_3 - \varepsilon_1} \qquad (S47)$$

The cell division time of the phagocytoser is obtained similarly but with $P_{nut} = 1$ and the cost of the trait being due to prey capture, engulfment, and internal digestion:

$$t_{d,phago} = \frac{N_{nutV}V_{pred}(1 + C_{trait})}{N_{AA,prey}v_{capadj}N_{value}} \qquad (S48)$$

Substituting the trait costs and $N_{value}$ yields:

$$t_{d,phago} = \frac{\alpha_1 + \alpha_2 + \alpha_3 + \alpha_4 + \beta_1 t_{d,anc}}{\alpha_5\left(1 - \frac{t_{dig}}{6t_{d,phago}}\right)} = \frac{\alpha_1 + \alpha_2 + \alpha_3 + \alpha_4 + \beta_1 t_{d,phago}}{\alpha_5 - \frac{\beta_2}{t_{d,phago}}} \qquad (S49)$$

$$\alpha_1 = N_{nutV}V_{pred} \qquad (S49a)$$

$$\alpha_2 = N_{nutV}V_{pred}C_{eng}\frac{1}{C_{budget}f_{budget}} \qquad (S49b)$$

$$\alpha_3 = N_{nutV}V_{pred}C_{cap}\frac{1}{C_{budget}} \qquad (S49c)$$

$$\alpha_4 = N_{nutV} V_{pred} C_{fv} \frac{1}{C_{budget} f_{budget}} \qquad (S49d)$$

$$\alpha_5 = N_{AA,prey} v_{capadj} \qquad (S49e)$$

$$\beta_1 = N_{nutV} V_{pred} \left( \frac{2N_{AA,prey}}{k_{prot} t_{dig}} N_{AA,enz} C_{AA} v_{capadj} \right) \frac{1}{C_{budget} f_{budget}} \qquad (S49f)$$

$$\beta_2 = N_{AA,prey} v_{capadj} \frac{t_{dig}}{6} \qquad (S49g)$$

The cell division time of the phagocytoser can be obtained from the alphas and betas by rewriting the equation:

$$t_{d,phago} = \frac{\alpha_1 + \alpha_2 + \alpha_3 + \alpha_4 + \beta_2}{\alpha_5 - \beta_1} \qquad (S50)$$

### Filters for data

The cells considered here suffer from several internal constraints that are not directly accounted for by the phagocytosis model itself. To make sure that the presented data don't include transgressions of these constraints, several filters are applied. These concern (1) the area limit imposed by the size of the food vacuole on the number of Sec translocases and nutrient transporters, (2) the limit imposed by the predator cell volume on the number of food vacuoles present simultaneously, and (3) the area limit imposed by the surface of the ancestral predator on the number of prey that is digested externally at the same time. In addition, the data is screened for fractional occupancies in the ancestor, $f_{occ,anc}$, and phagocytoser, $f_{occ,phago}$, that exceed 1.

#### *Food vacuole area limit*

The area occupied by Sec translocases and nutrient transporters on the food vacuole membrane is a function of their numbers. The number of Sec translocases is determined from:

$$N_{AA,prey} = \int_0^{t_{dig}} N_{enz}(t) k_{prot} dt = \int_0^{t_{dig}} N_{Sec} \frac{t}{t_{ins}} k_{prot} dt \qquad (S51)$$

Solving the integral yields:

$$N_{AA,prey} = \frac{N_{Sec} k_{prot} {t_{dig}}^2}{2 t_{ins}} \qquad (S52)$$

Giving the number of Sec translocases as:

$$N_{Sec} = \frac{2 N_{AA,prey} t_{ins}}{k_{prot} {t_{dig}}^2} \qquad (S53)$$

The maximum nutrient production rate for enzymes is $N_{enz} \frac{t_{dig}}{t_{ins}} k_{prot}$, this needs to matched by the nutrient transport rate, $N_{trans} k_{trans}$, yielding the number of nutrient transporters as:

$$N_{trans} = \frac{N_{enz} t_{dig} k_{prot}}{t_{ins} k_{trans}} \qquad (S54)$$

The area occupancy of the Sec translocases and nutrient transporters is:

$$A_{SecTrans} = \frac{A_{mp}}{f_{memprot}} (N_{Sec} + N_{trans}) \qquad (S55)$$

The fractional occupancy on the food vacuole is:

$$f_{occ,fv} = \frac{A_{SecTrans}}{A_{fv}} = \frac{A_{SecTrans}}{4\pi \left(\frac{3V_{prey}}{4\pi}\right)^{\frac{2}{3}}} \qquad (S56)$$

Datapoints for which $f_{occ,fv} > 1$ are removed from the analysis.

*Predator cell volume limit*

The number of food vacuoles that is present simultaneously is calculated from the digestion time and the prey capture rate:

$$N_{fv} = t_{dig} v_{cap} \qquad (S57)$$

The volume of the food vacuole is equal to that of the prey giving a total food vacuole volume of $N_{fv} V_{prey}$, which lead to a fractional volume occupancy of:

$$f_{occ,Vol} = \frac{t_{dig} v_{cap} V_{prey}}{V_{pred}} \qquad (S58)$$

Datapoints for which $f_{occ,Vol} > 1$ are removed from the analysis.

*Surface area limit for external digestion*

The number of prey cells being present simultaneously on the ancestor cell surface is calculated from the digestion time and the prey capture rate:

$$N_{prey} = t_{dig} v_{cap} \qquad (S59)$$

The area occupied by a single prey cell is $\pi {r_{prey}}^2$, yielding a fractional occupancy of prey cells on the predator cell surface of:

$$f_{occ,prey} = \frac{\pi {r_{prey}}^2 t_{dig} v_{cap}}{A_{pm}} \qquad (S60)$$

Datapoints for which $f_{occ,prey} > 1$ are removed from the analysis.

### Comparing phagocytoser to ancestor fitness

The fitness of the phagocytoser is calculated relative to that of the ancestor. This is done by a two-step method. First, the digestion time, $t_{dig}$, and the prey capture rate, $v_{cap}$, are varied and the cell division time and area occupancies are calculated for the ancestor and phagocytoser separately. Taking just the ancestor cell division times and area occupancies, fitness is calculated by comparing all parameter combinations to a single selected parameter combination. The same is performed for the phagocytoser. For both ancestor and phagocytoser the maximal fitness is found, and the cell division time and area occupancy associated with that maximal fitness are retrieved. In the second step, the fitness of the optimal phagocytoser is compared to the optimal ancestor for ranges of nutrient return probability, $P_{nut}$, and engulfment time, $t_{eng}$. For each combination of $P_{nut}$ and $t_{eng}$, step one is repeated and a fitness of phagocytoser is calculated relative to the ancestor.

### Phagocytosis intermediates

#### *No food vacuole fusion*

Phagocytosis in which food vacuoles don't fuse with the plasma membrane can be modeled by modifying the cell division equation for the phagocytoser (Eq. S50). The effect of the energy cost of the steady state food vacuole number, $\alpha_4$, needs to be removed and replaced by an effect similar to that of enzymes (Eq. S49f), yielding:

$$t_{d,phago} = \frac{\alpha_1 + \alpha_2 + \alpha_3 + \beta_2}{\alpha_5 - \beta_1 - \beta_3} \qquad (S61)$$

with

$$\beta_3 = N_{nutV} V_{pred} \left( A_{fv} C_{area} v_{capadj} \right) \frac{1}{C_{budget} f_{budget}} \qquad (S62)$$

$A_{fv}$ is the surface area of a single food vacuole and $C_{area}$ is the energy cost per unit membrane area (composed of lipids and membrane proteins).

#### *Low area availability on the food vacuole*

A food vacuole with low selectivity for Sec translocons and nutrient transporters can be modeled by adjusting the data selection criterion (see "*Food vacuole area limit*" section) for the surface area occupancy of Sec translocases and nutrient transporters, already included in the model, from 1 to 0.1.

#### *Large phagocytic cups*

Larger phagocytic cups are modeled by recalculating the energy cost and area occupancy for a phagocytic cup with thicker walls. The cost of the cup is given by:

$$C_{cup} = A_{cup} C_{area} + V_{cup} C_{plasm} \qquad (S63)$$

$C_{area}$ is the energy cost per unknit membrane and $C_{plasm}$ is the energy cost per unit cup plasm (see "*Energy cost of engulfment*" section). The area of the cup is given by:

$$A_{cup} = A_{prey} + 2\pi\left(r_{prey} + d_{cup}\right) r_{prey} + 2\pi\left(r_{prey} + d_{cup}\right)^2 - \pi\left(r_{prey} + d_{cup}\right)^2 \qquad (S64)$$

$A_{prey}$ is the surface area of the prey, $r_{prey}$ is the radius of the prey, and $d_{cup}$ is the thickness of the phagocytic cup. The volume of the cup is given by:

$$V_{cup} = \frac{1}{2}\frac{4\pi}{3}\left(r_{prey} + d_{cup}\right)^3 + \pi\left(r_{prey} + d_{cup}\right)^2 r_{prey} - V_{prey} \qquad (S65)$$

$V_{prey}$ is the prey volume. The surface area occupancy of the phagocytic cup is given by:

$$A_{occ,cup} = \pi\left(r_{prey} + d_{cup}\right)^2 \qquad (S66)$$

To determine the fitness consequences $d_{cup}$ is set to 1 μm and the area occupancy and energy cost so calculated are substituted into the fitness model.

### Protease turnover numbers

Protease turnover numbers were obtained from the BRENDA database, limiting the data to protein or peptide substrates. Turnover numbers vary by orders of magnitude and are listed in Table S3.

**Table S3: Protease turnover numbers**

| Enzyme | Organism | Substrate | Turnover Number ($s^{-1}$) | Reference |
|---|---|---|---|---|
| **Cytosol nonspecific dipeptidase** | Mus musculus | Ala-Gly | 43.3 | [26] |
| **Cytosol nonspecific dipeptidase** | Platyrrhini | Ala-Leu | 1190 | [27] |
| **Cytosol nonspecific dipeptidase** | Platyrrhini | Ala-Leu | 6510 | [27] |
| **Cytosol nonspecific dipeptidase** | Platyrrhini | Gly-Val | 7450 | [27] |
| **Cytosol nonspecific dipeptidase** | Platyrrhini | Gly-Phe | 8190 | [27] |
| **Cytosol nonspecific dipeptidase** | Platyrrhini | Gly-Ile | 10700 | [27] |
| **Cytosol nonspecific dipeptidase** | Platyrrhini | Gly-Leu | 10900 | [27] |
| **Cytosol nonspecific dipeptidase** | Platyrrhini | Gly-Phe | 20400 | [27] |
| **Cytosol nonspecific dipeptidase** | Platyrrhini | Gly-Val | 21400 | [27] |
| **Cytosol nonspecific dipeptidase** | Platyrrhini | Gly-Ile | 23000 | [27] |
| **Cytosol nonspecific dipeptidase** | Platyrrhini | Gly-Leu | 23100 | [27] |
| **Cytosol nonspecific dipeptidase** | Mus musculus | Ala-Ile | 33300 | [26] |
| **Endopeptidase Clp** | Escherichia coli | Oxidized insulin B-chain | 0.5 | [28] |
| **Endopeptidase Clp** | Escherichia coli | Insulin chain B | 0.35 | [29] |

| | | | | |
|---|---|---|---|---|
| **Endopeptidase Clp** | Escherichia coli | Glucagon | 0.5 | [28] |
| **Endopeptidase Clp** | Escherichia coli | Phe-Ala-Pro-His-Met-Ala-Leu-Val-Pro-Val | 13.3 | [28] |
| **Endopeptidase Lon** | Escherichia coli | Casein | 0.0333 | [30] |
| **Endopeptidase Lon** | Escherichia coli | Oxidized insulin B-chain | 0.333 | [30] |
| **Endopeptidase Lon** | Escherichia coli | Beta-galactosidase fragment 3-93 | 0.84 | [31] |
| **Endopeptidase Lon** | Escherichia coli | Bacteriophage lambda N-protein | 1 | [30] |
| **Leucyl aminopeptidase** | Bos taurus | Cys-Gly | 2.6 | [32] |
| **Leucyl aminopeptidase** | Salmonella enterica | Asp-Leu-Lys | 13 | [33] |
| **Leucyl aminopeptidase** | Bos taurus | Ser-Gly | 17 | [32] |
| **Leucyl aminopeptidase** | Salmonella enterica | Asp-Ile-Gly-Gly | 26 | [33] |
| **Leucyl aminopeptidase** | Bos taurus | Ser-Gly | 42 | [32] |
| **Leucyl aminopeptidase** | Bos taurus | Cys-Gly | 56.66 | [32] |
| **Leucyl aminopeptidase** | Bos taurus | Cys-Gly | 57 | [32] |
| **Leucyl aminopeptidase** | Bos taurus | Cys-Gly | 67 | [32] |
| **Leucyl aminopeptidase** | Bos taurus | Cys-Gly | 100 | [32] |
| **Leucyl aminopeptidase** | Bos taurus | Cys-Gly | 118 | [32] |
| **Leucyl aminopeptidase** | Bos taurus | Cys-Gly | 118.3 | [32] |
| **Leucyl aminopeptidase** | Bos taurus | Met-Gly | 135.8 | [32] |
| **Leucyl aminopeptidase** | Bos taurus | Met-Gly | 136 | [32] |
| **Leucyl aminopeptidase** | Bos taurus | Leu-Gly | 191.7 | [32] |
| **Leucyl aminopeptidase** | Bos taurus | Leu-Gly | 192 | [32] |
| **Leucyl aminopeptidase** | Salmonella enterica | Leu-Tyr | 351 | [33] |
| **Leucyl aminopeptidase** | Bos taurus | Leu-Gly | 400 | [32] |

| | | | | |
|---|---|---|---|---|
| **Leucyl aminopeptidase** | Salmonella enterica | Asp-His | 418 | [33] |
| **Leucyl aminopeptidase** | Bos taurus | Met-Gly | 468 | [32] |
| **Leucyl aminopeptidase** | Bos taurus | Met-Gly | 486.3 | [32] |
| **Leucyl aminopeptidase** | Salmonella enterica | Asp-Phe | 500 | [33] |
| **Leucyl aminopeptidase** | Salmonella enterica | Asp-Tyr | 500 | [33] |
| **Leucyl aminopeptidase** | Salmonella enterica | Asp-Leu | 666 | [33] |
| **Leucyl aminopeptidase** | Bos taurus | Leu-Gly | 675 | [32] |
| **Leucyl aminopeptidase** | Salmonella enterica | Asp-Gln | 761 | [33] |
| **Leucyl aminopeptidase** | Salmonella enterica | Glu-Leu | 866 | [33] |
| **Leucyl aminopeptidase** | Salmonella enterica | Leu-Leu | 918 | [33] |
| **Leucyl aminopeptidase** | Salmonella enterica | Asp-Leu-Gly | 950 | [33] |
| **Leucyl aminopeptidase** | Bos taurus | Met-Gly | 988 | [32] |
| **Leucyl aminopeptidase** | Bos taurus | Met-Gly | 988.3 | [32] |
| **Leucyl aminopeptidase** | Solanum lycopersicum | Arg-Gly-Asp | 3781 | [34] |
| **Leucyl aminopeptidase** | Solanum lycopersicum | Gly-Leu | 5140 | [34] |
| **Leucyl aminopeptidase** | Solanum lycopersicum | Gly-Leu | 5141 | [34] |
| **Leucyl aminopeptidase** | Solanum lycopersicum | Arg-Gly-Phe | 6070 | [34] |
| **Leucyl aminopeptidase** | Solanum lycopersicum | Arg-Gly-Phe | 6073 | [34] |
| **Leucyl aminopeptidase** | Solanum lycopersicum | Gly-Gly | 10860 | [34] |
| **Leucyl aminopeptidase** | Solanum lycopersicum | Gly-Gly | 10900 | [34] |
| **Leucyl aminopeptidase** | Solanum lycopersicum | Arg-Gly | 13500 | [34] |
| **Leucyl aminopeptidase** | Solanum lycopersicum | Arg-Gly | 13520 | [34] |
| **Leucyl aminopeptidase** | Solanum lycopersicum | Leu-Gly-Gly | 13960 | [34] |

| | | | | |
|---|---|---|---|---|
| **Leucyl aminopeptidase** | Solanum lycopersicum | Leu-Gly-Gly | 14000 | [34] |
| **Leucyl aminopeptidase** | Solanum lycopersicum | Leu-Gly | 14100 | [34] |
| **Leucyl aminopeptidase** | Solanum lycopersicum | Leu-Gly | 14400 | [34] |
| **Leucyl aminopeptidase** | Solanum lycopersicum | Leu-Gly | 14440 | [34] |
| **Leucyl aminopeptidase** | Solanum lycopersicum | Arg-Leu | 19380 | [34] |
| **Leucyl aminopeptidase** | Solanum lycopersicum | Arg-Leu | 19400 | [34] |
| **Leucyl aminopeptidase** | Solanum lycopersicum | Leu-Leu | 34490 | [34] |
| **Leucyl aminopeptidase** | Solanum lycopersicum | Leu-Leu | 34500 | [34] |
| **PepB aminopeptidase** | Salmonella enterica | Leu-Leu | 918 | [33] |
| **PepB aminopeptidase** | Salmonella enterica | Asp-Leu-Gly | 950 | [33] |
| **PepB aminopeptidase** | Salmonella enterica | Glu-Leu | 866 | [33] |
| **Tripeptide aminopeptidase** | Homo sapiens | Arg-Ser-Arg | 0.56 | [35] |
| **Tripeptide aminopeptidase** | Homo sapiens | Arg-Ala-Arg | 5.3 | [35] |
| **Tripeptide aminopeptidase** | Sus scrofa | Gly-Gly-Gly | 2030 | [36] |

### Parameters

The various parameters used in the phagocytosis model and the supporting simulations are described in the supplementary information and a subset is listed in Table S4.

**Table S4: Parameter values.**

| Parameter | Value | Unit | References and Comments |
|---|---|---|---|
| $\boldsymbol{N_{nutV}}$ | 2x10$^9$ | N/A | [37], p. 223 |
| $\boldsymbol{V_{pred}}$ | 1000 | μm$^3$ | Typical volume of a eukaryote[38]. |
| $\boldsymbol{V_{prey}}$ | 1 | μm$^3$ | Typical volume of a prokaryote[38]. |
| $\boldsymbol{c_{prey}}$ | 0.0001 | μm$^{-3}$ | Prey concentration. |
| $\boldsymbol{N_{AA,prey}}$ | 2x10$^9$ | N/A | [37], p. 223 |
| $\boldsymbol{N_{AA,enz}}$ | 350 | N/A | Typical protein length. |
| $\boldsymbol{C_{AA}}$ | 29 | ATP | Energetic cost of an average amino acid[25]. |
| $\boldsymbol{P_{nut}}$ | 0.88 | N/A | Probability for nutrients of reaching the predator. This work. |

| | | | |
|---|---|---|---|
| $P_{enz}$ | 0.067 | N/A | Probability for enzymes of reaching a single prey cell. This work. |
| $A_{phago}$ | 1.67 | $\mu m^2$ | Area occupancy of phagocytic cup at maximum extent for a prey volume of 1 $\mu m^3$. This work. |
| $C_{axo}$ | $1.83 \times 10^{11}$ | ATP | Energy cost of an individual axopod. This work. |
| $r_{axo}$ | 0.14 | µm | Radius of an axopod. This work, see Table S1 and Figure S1A. |
| $A_{mp}$ | $10^{-5}$ | $\mu m^2$ | Area occupancy of individual Sec translocases and nutrient transporters. |
| $k_{prot}$ | 100 | $s^{-1}$ | Amino acid production rate by enzymes. See Table S3. |
| $k_{trans}$ | 10 | $s^{-1}$ | Nutrient transporter rate. Value unclear. Various sugar transporters have a range of 20-300 molecules $s^{-1}$ (p. 224 in ref[37]). An in vitro estimate for aspartate transport[39] is 0.01 $s^{-1}$. |
| $t_{ins}$ | 20 | s | Time of enzyme secretion by a Sec translocase[1,40]. |
| $f_{memprot}$ | 0.4 | N/A | Fraction of membrane area that is occupied by transmembrane proteins[41]. |
| $f_{budget}$ | 1.5 | N/A | Growth modifier to the cell budget. This work. |
| $C_{area}$ | $1.45 \times 10^{9}$ | ATP $\mu m^{-2}$ | Energy cost of membrane per unit area (includes lipids and proteins)[11]. |

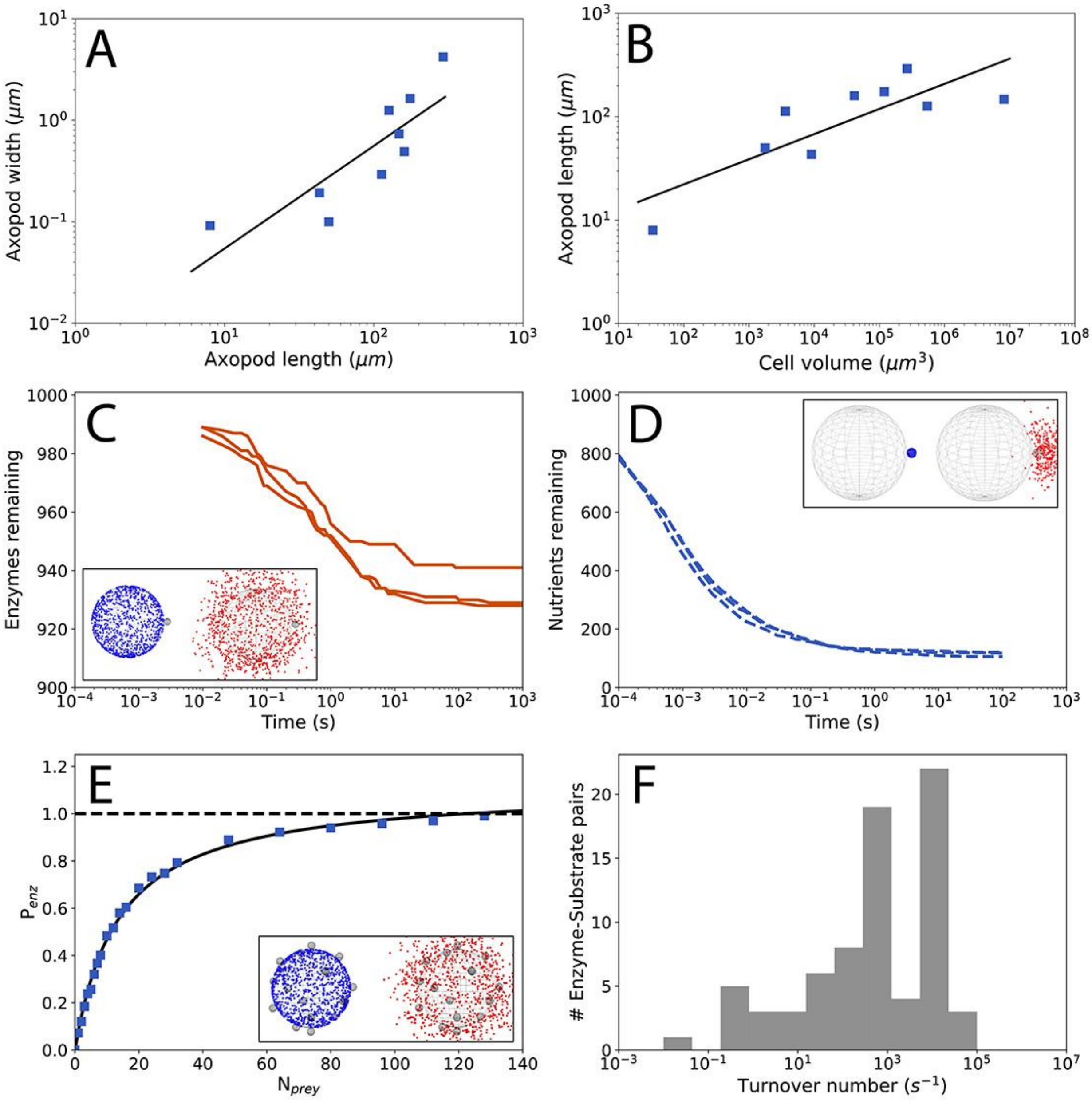


**Figure S1:** A) Axopod lengths and widths for heliozoans[3–9]. Equation for the regression: $y = 0.0053\, x^{1.00}$. B) Heliozoan cell volumes and axopod lengths[3–9]. Equation for the regression: $y = 7.24\, x^{0.24}$. C) Smoldyn simulation results of enzyme finding prey, leading to a probability of finding prey, $P_{enz} = 0.067$. Inset: simulation setup with predator surface-attached enzymes in blue on the left, and free enzymes in red on the right. D) Smoldyn simulation results of nutrient finding the predator, leading to a probability of finding the predator, $P_{nut} = 0.88$. Inset: simulation setup with prey surface-attached nutrients in blue on the left, and free nutrients in red on the right. E) Smoldyn simulation results of enzyme finding prey, with varying prey number. Inset: simulation setup with predator surface-attached enzymes in blue on the left, and free enzymes in red on the right. F) Distribution of protease turnover numbers of enzyme-substrate pairs (Table S3).

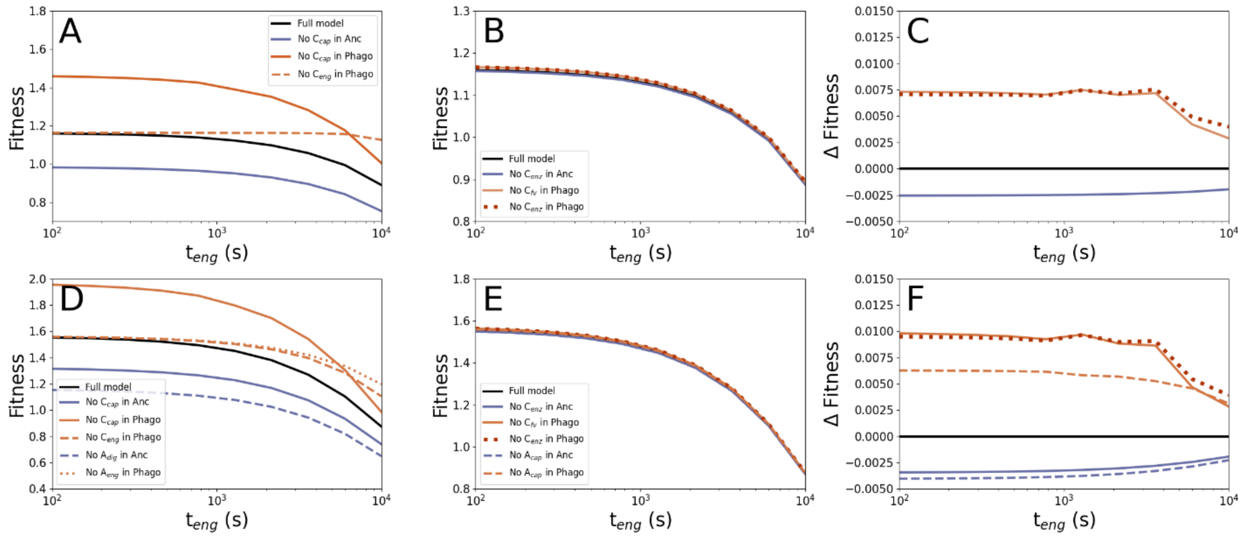


**Figure S2: Analysis of fitness penalties associated with energy and surface area occupancy costs.** A-C) Fitness calculated using cell division times only. A) The full model compared to models lacking one of the larger fitness penalties. B) Comparison of full model to models lacking the smaller fitness penalties. C) Fitness difference by subtracting the full model fitness from the fitness of the models lacking the smaller fitness penalties (legend as in B). D-F) Fitness calculated using cell division times and surface area occupancies. D) The full model compared to models lacking one of the larger fitness penalties. E) Comparison of full model to models lacking one of the smaller fitness penalties. F) Fitness difference by subtracting the full model fitness from the fitness of the models lacking the smaller fitness penalties (legend as in E).

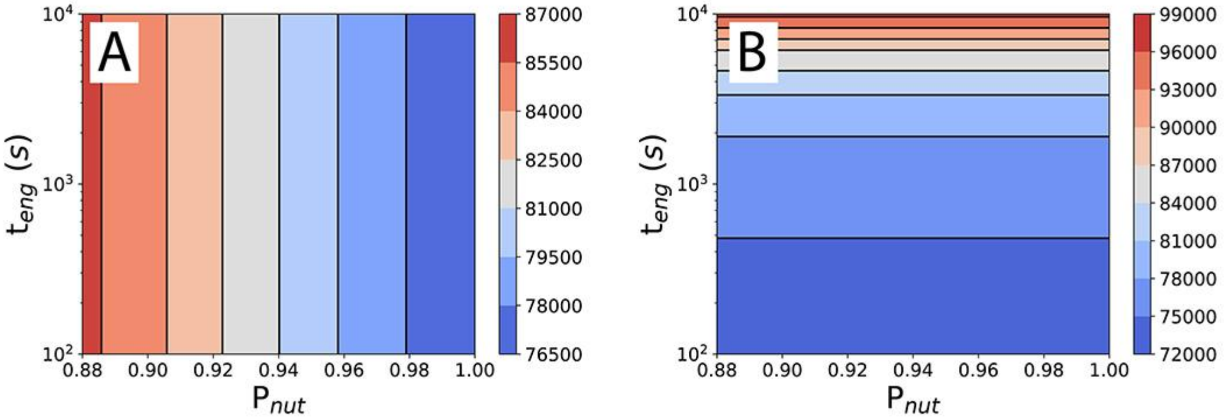


**Figure S3: Cell division times of the ancestor (A) and phagocytoser (B).** Cell division times in seconds. Related to fitness calculations in Figure 2F and 2G.

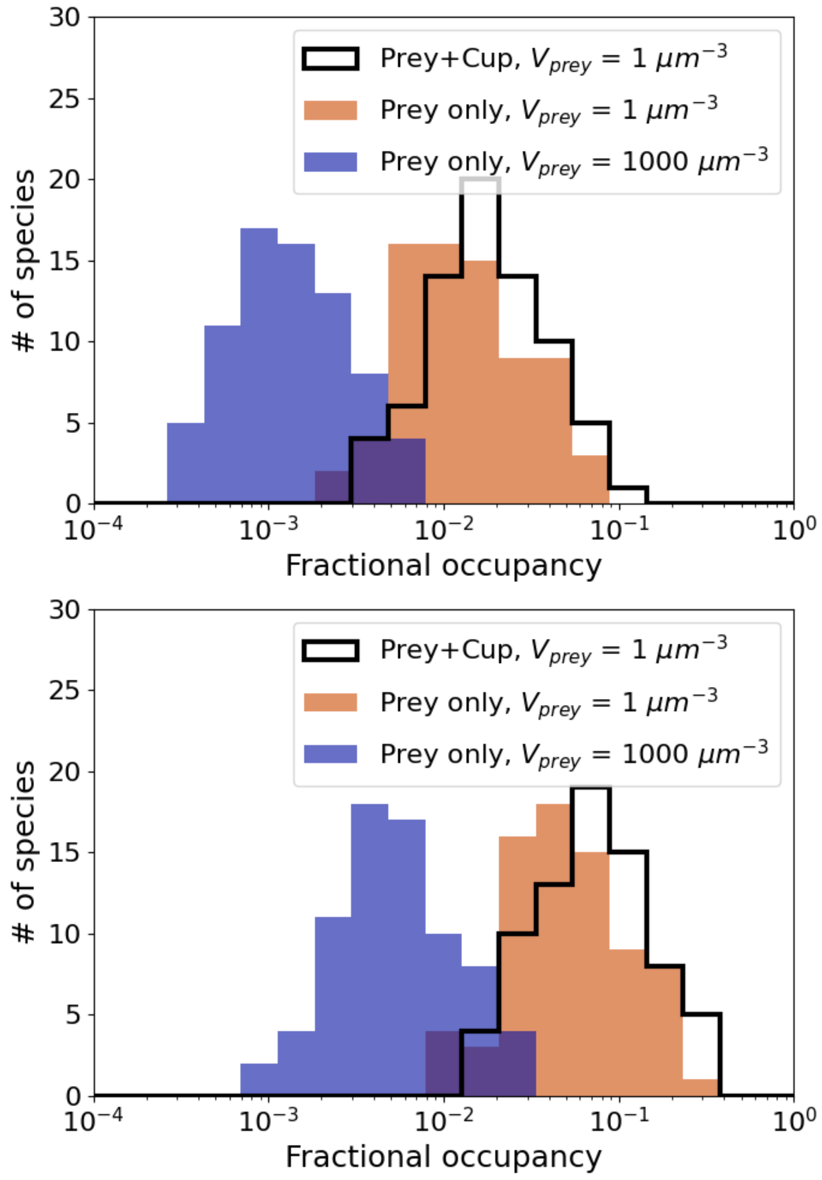


**Figure S4: Predictions for surface occupancy of phagocytosis on the cell surface of ciliate species.** Top panel, fractional occupancies calculated assuming biomass conversion efficiency of 100%. Bottom panel, fractional occupancies calculated assuming biomass conversion efficiency of 25%. Calculated from ciliate cell volumes and cell division times obtained from[42]. Fractional occupancy in the area occupied by phagocytic cups divided by the surface area of the ciliate (assuming that the ciliates are spheres, relaxing this assumption would push the graphs toward even lower fractional occupancy). For the black line (Prey+Cup), a phagocytic cup area equal to $A_{phago}/2$ is used, with the total area depending on the engulfment time (180 s) and the rate of prey capture (calculated from the prey cell volume, ciliate cell volume, and the ciliate cell division time). For the orange and blue distributions the fractional occupancy is obtained in the same way, except that the area occupancy per prey cell is calculated using $\frac{\pi}{2}\left(\frac{3V_{prey}}{4\pi}\right)^{\frac{2}{3}}$, which ignores the contribution to the area by the walls of the phagocytic cup.
